# A simple model for detailed visual cortex maps predicts fixed hypercolumn sizes

**DOI:** 10.1101/2020.09.01.277319

**Authors:** Marvin Weigand, Hermann Cuntz

## Abstract

Orientation hypercolumns in the visual cortex are delimited by the repeating pinwheel patterns of orientation selective neurons. We design a generative model for visual cortex maps that reproduces such orientation hypercolumns as well as ocular dominance maps while preserving retinotopy. The model uses a neural placement method based on t–distributed stochastic neighbour embedding (t–SNE) to create maps that order common features in the connectivity matrix of the circuit. We find that, in our model, hypercolumns generally appear with fixed cell numbers independently of the overall network size. These results would suggest that existing differences in absolute pinwheel densities are a consequence of variations in neuronal density. Indeed, available measurements in the visual cortex indicate that pinwheels consist of a constant number of ∼30, 000 neurons. Our model is able to reproduce a large number of characteristic properties known for visual cortex maps. We provide the corresponding software in our `MAPStoolbox` for *Matlab*.

**In brief:** We present a generative model that predicts visual map structures in the brain and a large number of their characteristic properties; a neural placement method for any given connectivity matrix.

**Highlights:**

- Generative model with retinotopy, orientation preference and ocular dominance.
- Prediction of constant neuronal numbers per orientation hypercolumn.
- Curated data shows constant ∼30, 000 neurons per pinwheel across species.
- Simple explanation for constant pinwheel and orientation hypercolumn ratios.
- Precise prediction of ∼80% nearest neighbour singularities with opposing polarity.
- Model asymptotically approaches realistic normalised pinwheel densities.
- Small brains with < ∼300 potential pinwheels exhibit salt-and-pepper maps.
- Different map phenotypes can exist even for similar connectivity.

## Introduction

Cortical neurons across layers typically respond to the same representational feature, forming a columnar arrangement (Mountcastle, 1997; Kaas, 2012). Since feature preferences of neurons change continuously along the cortical surface rather than in discrete steps, a natural subdivision into discrete columns is, however, not generally obvious. On the other hand, continuously repeating patterns encompassing a complete set of values in one feature dimension can more easily be distinguished; these form cortical hypercolumns that divide the innately continuous cortical maps into maps of discrete cortical patches (Horton and Adams, 2005). In particular, orientation selectivity in the primary visual cortex has been used extensively to study cortical hypercolumns (Hubel and Wiesel, 1974; Blasdel and Salama, 1986; Bonhoeffer et al., 1995; Ohki et al., 2006; White and Fitzpatrick, 2007). However, the biological significance of this type of anatomical blueprint remains elusive.

A large diversity of cortical map models has allowed for an increasingly quantitative understanding of the organisation of hypercolumns in visual cortex. Because of the inherent dependence of hypercolumn structure on visual input (Constantine-Paton and Law, 1978; DeBruyn and Casagrande, 1981; Sengpiel et al., 1996; Sharma et al., 2000; White et al., 2001), activity dependent mechanisms have been linked to their self-organised formation. Accordingly, many existing models rely on a predefined grid of neurons that refine their feature preferences iteratively based on a given input (see Erwin et al., 1995; Swindale, 1996; Goodhill, 2007, for review). While some models implement a nerve net with firing neurons and Hebbian learning rules (von der Malsburg, 1973, 1979; Linsker, 1986a,b,c; Miller et al., 1989; Ernst et al., 2001; Stevens et al., 2013), others omit the biological details and predict maps for a given set of feature vectors representing the retinotopic, ocular dominance, orientation and direction preference of neurons in the visual cortex (Durbin and Mitchison, 1990; Obermayer et al., 1990; Swindale and Bauer, 1998). The more abstract models particularly based on elastic net (EN) (Durbin and Willshaw, 1987) and Kohonen map algorithms (Kohonen, 1982), are able to reproduce many characteristics of visual cortex maps (Erwin et al., 1995; Swindale, 1996).

Assuming that visual cortex maps are formed by activity-dependent principles, some of these characteristics could be linked with function of the neural network or connectivity. In particular, differences across species are useful to identify such functional requirements for a given anatomical formation. Most strikingly, orientation selective neurons in rodents are scattered in a salt-and-pepper pattern, but are organised according to their preferred orientation in the aforementioned pinwheel-like arrangements in other mammals forming discrete orientation hypercolumns (Kaschube, 2014). Intuitively, such morphological differences could be a consequence of fundamental architectural differences in the neural circuit design of these animals. Tracing experiments in cats (Gilbert and Wiesel, 1989), tree shrews (Fitzpatrick, 1996; Bosking et al., 1997) and macaques (Malach et al., 1993) showed that connections between neurons are formed preferably between neurons of similar orientation preference which was recently confirmed at the synapse level for tree shrews (Zhang et al., 2018). Based on such a like-to-like connectivity, pinwheel arrangements similar as those found in these animals were predicted by wiring optimisation principles whereas salt-and-pepper patterns were predicted for random connectivities (Koulakov and Chklovskii, 2001). However, neurons in the salt-and-pepper cortex of rodents exhibit a similar bias for connections with other neurons of similar orientation preference (Ko et al., 2011, 2013, 2014; Lee et al., 2016). This would exclude that the salt-and-pepper map in this case is a consequence of a random connectivity between orientation selective neurons despite the distinct morphological phenotypes.

We have previously designed a class of novel simple models that do not depend on the activity of neurons or on a predefined grid of model neurons (Weigand et al., 2017). Instead, the neuronal map layout is predicted from a given connectivity that may or may not have been shaped by activity dependent principles. The model is based on dimensionality reduction methods and establishes relative neuronal positions for arbitrary connectivities. This approach is simple to use and helps to elucidate the links between the details of neural circuits and the corresponding anatomical structures allowing for a clear functional interpretation while remaining at a phenomenological non-biological level of the implementation. Using these models, we have shown a phase transition from single pinwheels to seemingly unstructured salt-and-pepper maps by lowering the overall number of neurons without changing the specific selectivity of the connections. This could explain experimentally observed structured maps in larger animals such as cats and monkeys without assuming differences in the underlying connectivity.

Here, we use our models to better understand the detailed relations between the different features of visual cortex maps in the case of pinwheel arrangements. In particular, we focus on the density of pinwheels and the orientation hypercolumn area that were shown to have a constant relation in mammals of different orders (Kaschube et al., 2010). Interestingly, the size of orientation hypercolumns and thus their absolute density both vary widely (**Table S1**) (Yicong et al., 2012). The difference between absolute and normalised orientation hypercolumn densities could therefore be explained either by variations in the absolute number of neurons per orientation hypercolumn, by variations in the density of neurons with a constant number of neurons per orientation hypercolumn, or by a combination of both.

## Material and methods

All calculations and simulations were performed using custom code in *Matlab* (Mathworks) and were executed using the Neuroscience Gateway (Sivagnanam et al., 2013). We will make our code and data publicly available after the manuscript has been published.

### Optimal neuronal placement

Based on the idea that connected neurons should be located near each other to save overall wiring, we have previously introduced a model to predict some aspects of cortical maps using multi-dimensional scaling (MDS) (Weigand et al., 2017). Connection dissimilarities between neurons were used to predict relative neuronal positions such that the divergence between spatial distances and transformed connection dissimilarities was minimal. Placing neurons by using their connection dissimilarities to estimate their spatial distances is supported by experimental findings that relate a higher connection dissimilarity to a lower number of connections (Song et al., 2014) and a lower number of connections to a higher distance between cortical areas (Ercsey-Ravasz et al., 2013). It is intuitive that such a placement of neurons saves wiring length, since neurons that share a higher number of connections are placed closer to one another than neurons that share fewer connections. Therefore, using dimension reduction methods to place neural structures based on their connection dissimilarity should lead to a global reduction of the wiring cost. Placing cortical areas with MDS to reflect connection dissimilarities in the resulting spatial distances between areas consequently reproduced the general functional layout of the cortex (Young, 1992; Young et al., 1995; Song et al., 2014). A similar relationship may hold for individual neurons instead of cortical areas. Accordingly, using ordinal MDS (oMDS) enabled us to predict single pinwheels for a binary model connectivity that depended on the similarity of the orientation preference between neurons (Weigand et al., 2017). Briefly, Jaccard distances (JDs) and shortest path lengths (SPLs) were combined to calculate the connection dissimilarity of neurons according to their connectivity in the circuit. The resulting dissimilarity matrix was then subjected to oMDS to obtain the relative placement of neurons that best fit the distances in the matrix. This procedure enabled us to investigate how neuronal connectivity could result in the particular layouts observed in biology. In principle, the described method is compatible with any developmental mechanism that minimises the wiring length. However, the movement of neurons during the optimisation procedure is not corresponding to any realistic developmental mechanism.

In the present study, we modified this neural placement method by using cosine distances (CDs) instead of a combination of JD and SPL to calculate the connection dissimilarities and t–Distributed Stochastic Neighbour Embedding (t–SNE) (van der Maaten and Hinton, 2008) instead of oMDS to place the neurons in two-dimensional space based on the calculated connection dissimilarities. Using the CD as a dissimilarity measurement, the connection dissimilarities *δ*_*ij*_ between neurons *i* and *j* were defined as follows:

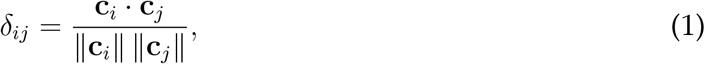

where **c**_*i*_ and **c**_*j*_ were the connection vectors of neurons *i* and *j*.

To calculate an optimal neuronal placement, the matrix **Δ** containing all pairwise connection dissimilarities *δ*_*ij*_ served as input for the t–SNE procedure from (van der Maaten and Hinton, 2008). To find the neuronal positions **Y** a cost function *C*, which was the Kullback-Leibler (KL) distance between pairwise probabilities *p*_*ij*_ and *q*_*ij*_, was minimised using t–SNE. This cost function was defined as follows:

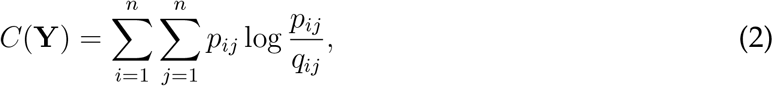

where *p*_*ij*_ was the probability that neuron *i* and *j* are neighbours based on their connection dissimilarity *δ*_*ij*_ and *q*_*ij*_ was the respective probability in the neuronal arrangement **Y** that depended on the Euclidean distance *d*_*ij*_ between neuron *i* and *j*. The pairwise probabilities 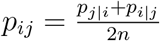 were defined by the symmetrised conditional probabilities

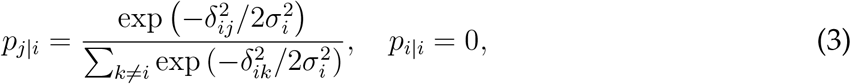

where *σ*_*i*_ was the variance of a Gaussian centered at neuron *i*. The variance of the Gaussians *σ*_*i*_ was calculated separately for every data point such that the perplexity 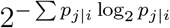 was constant. The probabilities *q*_*ij*_ were defined as follows

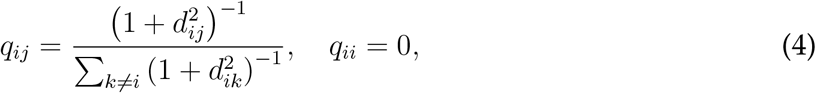

The cost function *C* was finally minimised by using gradient descent where the gradient

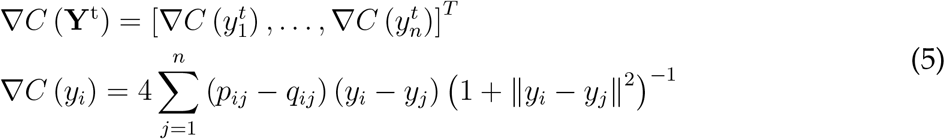

was used in each iteration *t* of the algorithm (see van der Maaten and Hinton, 2008, for derivation of the gradient and further details of the optimisation procedure).

We chose t–SNE over oMDS in this study specifically because it enabled us to calculate maps for feature dependent connectivities that are based on multiple features. Although oMDS leads to similar results for our simple hypercolumn model (**Figure S3**), it fails to separate connectivities depending on multiple features for our complete visual cortex model (**Figure S6**). A proof of concept of the method is shown in **Figure S1** (see Figure 1A in Weigand et al., 2017). Here, both methods are confronted with the same benchmark procedure as described previously (Weigand et al., 2017). t–SNE delivered comparable results but degenerate solutions appeared occasionally, which were discarded (**Figure S2A**). The error between the alignment of recovered and original positions was slightly higher than when using oMDS (4.59% compared to 2.09% positional deviation compared to the side length of the unit square, mean of 51 trials). In contrast, wiring length was slightly shorter than with oMDS (34.87% vs. 35.1% of the cable required for a random arrangement and 96.81% vs. 97.91% of the cable required for the original arrangement, mean of 51 trials). It is worth noting that these results can be explained with the tendency of the model neurons to be clustered when using t–SNE leading to arrangements that match the original arrangement less well but have a shorter overall wiring length (**Figure S1**). For all of our results we used t–SNE with the given standard parameters (perplexity was set to 30 and number of iterations to 1, 000).

**Figure 1.**
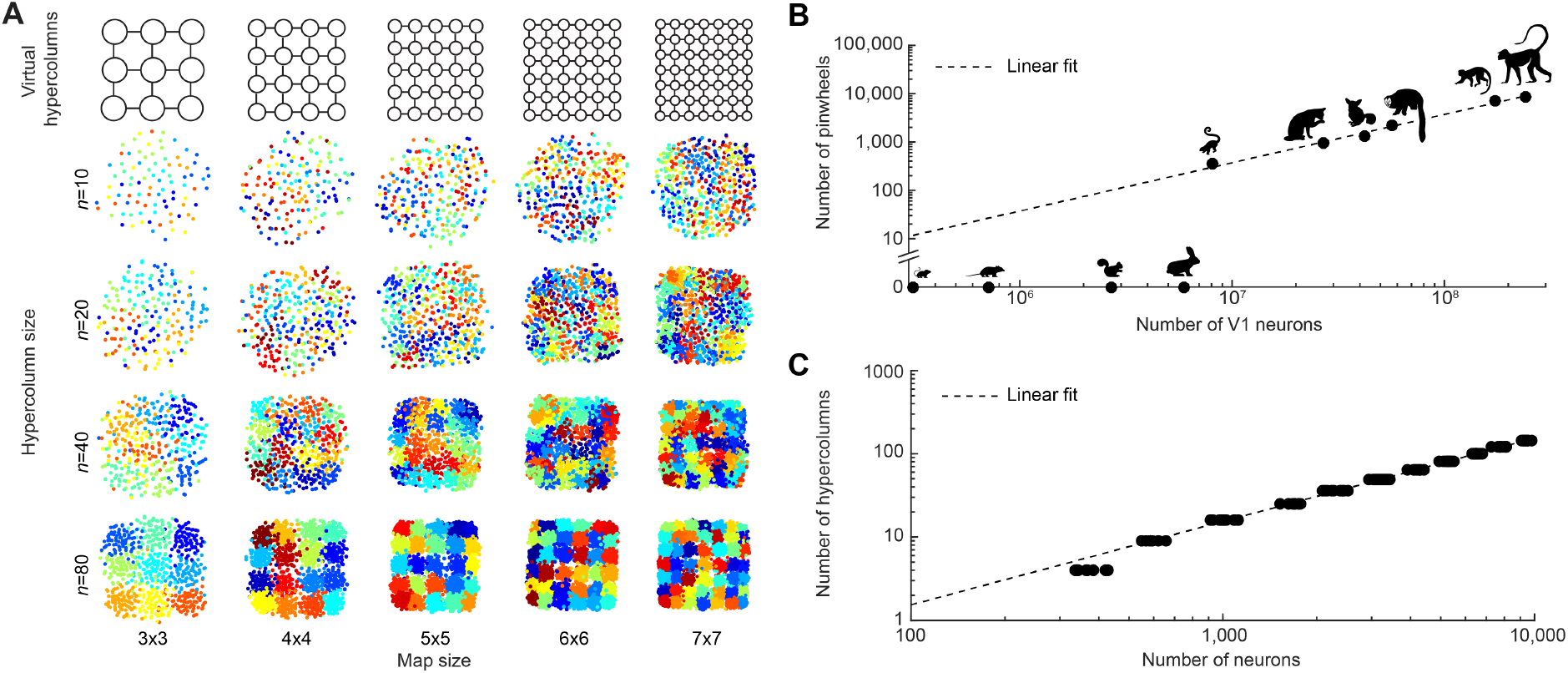
Map structure depends on neuron numbers per hypercolumn but not on overall map size. **A**, Individual maps of different sizes (horizontal) and different numbers of neurons per hypercolumn (vertical). Neurons (dots in the maps) assigned to the same hypercolumn in the underlying connectivity matrix are assigned to one random colour. **B**, The number of pinwheels plotted against the number of neurons in different species (from left to right: mouse, rat, squirrel, rabbit, tree shrew, cat, galago, owl monkey, squirrel monkey and macaque). The slope of the linear fit through the origin (black dashed line) for the species with pinwheels indicates a constant number of approximately 26, 840 neurons per pinwheel (95% confidence interval: 24, 800–28, 880, *R*^2^ = 0.9907). **C**, Number of hypercolumns and neurons at which neuronal map structure appears in our model defined by a scatter value of 0.5 (see **Materials and Methods**). Linear fit through the origin is indicated with black dashed line (*R*^2^ = 0.9975).

### Virtual hypercolumn model (grid model)

To model a simplified case of a cortical map where neurons cluster into hypercolumns, we assigned neurons equally to virtual hypercolumns on a quadratic grid of side lengths from 2 to 12 thus containing between 4 and 144 hypercolumns. Based on this arrangement we created a connectivity, which we used to predict the neuronal locations by the method described above. The connection between every pair of neurons was randomly determined using a connection probability that exponentially decayed with the Euclidean distance of the virtual hypercolumns to which both neurons were assigned, which is in accordance with empirically derived connectivity rules (Ercsey-Ravasz et al., 2013; Horvát et al., 2016). Accordingly, the connection probability *p*_*ij*_ between neurons *i* and *j* was set as follows:

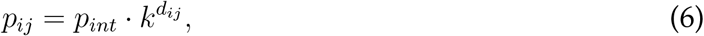

where *p*_*int*_ was the connection probability for neurons *i* and *j* assigned to the same virtual hypercolumn and *k* determined how steeply the connection probability decayed with the Euclidean distance *d*_*ij*_. We set *p*_*int*_ = 0.25 and *k* = 0.5 for all calculations.

To measure the amount of structure in the resulting maps, we calculated the scatter *s*. The scatter *s* describes the average relative number of neurons lying inside the space of any given hypercolumn but being associated with another hypercolumn:

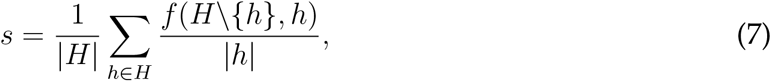

where *H* was the set of all virtual hypercolumns, *h* a set of neurons assigned to one virtual hypercolumn and *f* (*A, B*) was the function which returned the number of neurons of all hypercolumns in *A* lying inside the convex hull of all neurons of *B*.

We performed neuronal placement calculations for a fixed number of neurons but different map sizes (the resulting maps are partially shown in **Figure 1A**) and calculated the amount of structure given by *s* (**Figure S4**). Some of the resulting maps of the calculations for **Figure 1A** and **Figure S4** were distorted (**Figure S2B**) and were left out of the analysis in **Figure S4**. Using *s* as a measure for map structure, we were able to calculate maps of similar structure (which were used for the fit in **Figure 1C**). To obtain these similarly structured maps, we started using 10 neurons per virtual hypercolumn and increased the number of neurons per virtual hypercolumn until *s ≤* 0.5. This procedure was repeated 10 times for each map size.

### Visual cortex model

Using our placement method with a neuronal connectivity based on orientation and retinotopic preferences of neurons as found in the visual cortex of mammals we were able to model maps that contained structured orientation hypercolumns (**Figure 2C**). Since neurons in the primary visual cortex are preferably connected to neurons with similar feature preferences (Gilbert and Wiesel, 1989; Malach et al., 1993; Fitzpatrick, 1996; Bosking et al., 1997; Ko et al., 2011, 2013, 2014; Lee et al., 2016; Zhang et al., 2018), the input connectivity for the neuronal placement had to be created accordingly. We obtained the corresponding connectivity by first assigning to the *N* model neurons unique retinotopic preferences on a grid *x, y* ∈ [0, 1]. Additionally, we randomly assigned to each neuron one out of 100 orientation preferences *θ* ∈ Θ from 0 to *π* (Θ = [0, *π*], |Θ| = 100) with an equal spacing. We ensured that an equal number of neurons represented each of the 100 orientation preferences. Retinotopic preferences were defined as relative sizes between 0 and 1 corresponding to the minimum and maximum retinotopic coordinate of a V1 segment. Depending on the number of neurons in V1, the receptive field size *λ* varies (**Figure 2A**). Accordingly, the connection probability regarding the retinotopic preference between neurons *i* and *j* was made to depend on *λ* according to

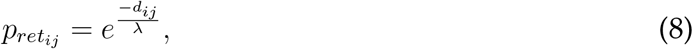

where *d*_*ij*_ was the Euclidean distance between the retinotopic preferences of neurons *i* and *j* (instances of connection function shown in **Figure S8A**, *left*). Similarly as for the simple grid model, we used an exponential relationship between the difference in retinotopic preference and connection probability, since retinotopy is continuously mapped along the cortical surface where connection probability decays exponentially with distance (Ercsey-Ravasz et al., 2013; Horvát et al., 2016).

**Figure 2.**
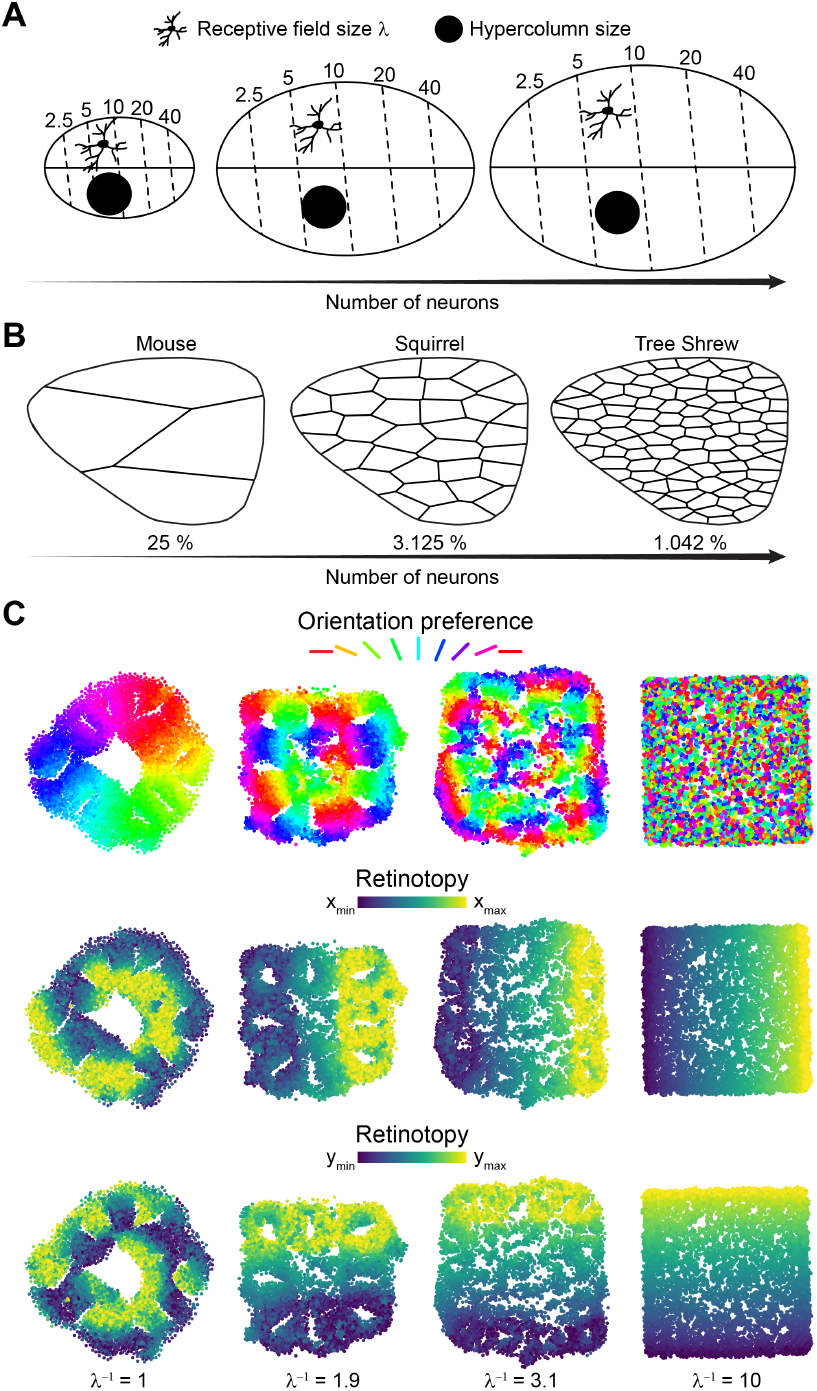
Relationship between number of neurons, receptive field size and map structure. **A**, Visual cortex size defined by its number of neurons influences the receptive field size *λ* and the proportion of a hypercolumn in the receptive field as indicated by the size relationships of a neuron and the space taken up by a hypercolumn (black circle) in the illustration based on Figure 4 in Kaas (2000) and Figure 1 in Elston et al. (1996). The numbers correspond to the visual angle of the dashed isoeccentricity lines and the midline to the horizontal meridian in this schematic illustration of topographic maps. **B**, Orientation hypercolumn occupancy and proportions of the visual cortex (given in percent) for respective predicted numbers of orientation hypercolumns (see **Materials and Methods**). Here, this is illustrated by a Voronoi diagram on regularly distributed points (not shown) representing the centers of orientation hypercolumns in a schematic drawing of V1 for three different species. **C**, OP maps (*top*) and retinotopic preference maps (*bottom*) produced by neuronal placement using a connectivity based on orientation preference and retinotopy. Only the receptive field size *λ* of neurons was varied as indicated. The number of neurons in all maps was kept constant at 6, 400 neurons (dots in the maps). All other parameters that determine the connection probability between neurons were also fixed for the shown model results (see **Materials and Methods**).

**Figure 3.**
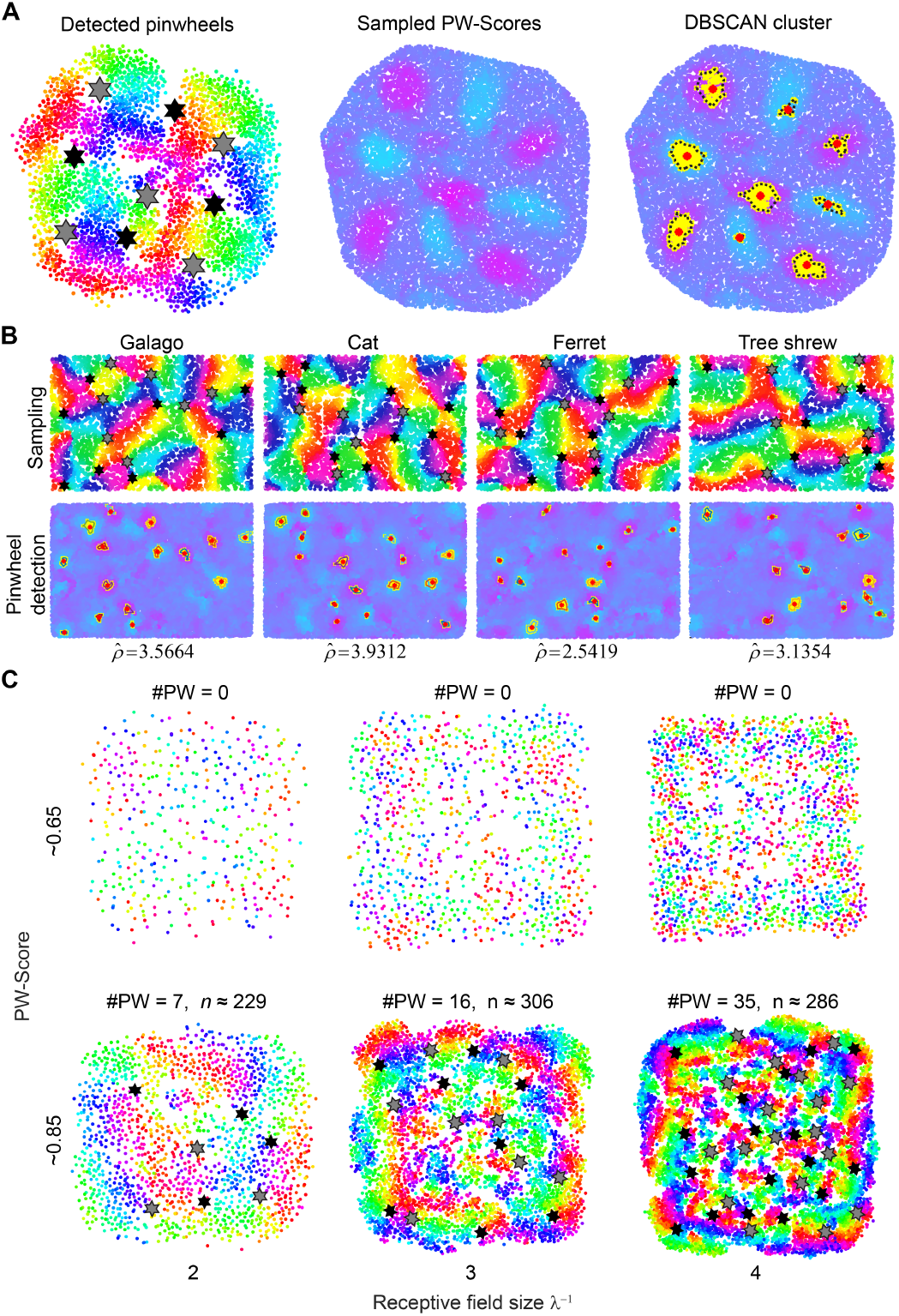
Detecting pinwheels in visual cortex maps of mammals and in the visual cortex model suggests constant orientation hypercolumn sizes. **A**, Detection of pinwheels in the modeled orientation preference maps. (*left*) Representative example of pinwheel detection for a map with 3, 600 neurons. Clockwise pinwheels are marked by a gray star and counter-clockwise pinwheels by a black star. (*middle*) Sampling of a score quantifying the *pinwheelness* at 20, 000 random sampling points in the OP map. The sampled score is colour coded where magenta indicates a perfect clockwise and cyan a perfect counter-clockwise pinwheel (see **Material and Methods**). (*right*) Pinwheels (red stars) are the centers of clusters (black broken lines) obtained from sampled score values over a threshold of 0.6 (yellow crosses). **B**, (*top*) Maps sampled from raster images (taken from Figure 2 in Kaschube, 2014) of OP maps in galago, cat, ferret and tree shrew to test our pinwheel detection method (see **Material and Methods**). The detected pinwheels are shown in the sampled maps as either gray or black stars corresponding to clock- and counterclockwise pinwheels. (*bottom*) Sampled pinwheel scores and pinwheel detection as in **A** (*right*). Calculated normalised pinwheel densities 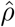 as for our model results (**Figure 4**) **are shown below. C**, Sample instances of OP maps generated by the visual cortex model for different representative numbers of neurons *N* and receptive field sizes *λ*. The maps were selected to have similar PW-scores in each row. Number of pinwheels (#PW) and number of neurons per pinwheel *n* are shown. Gray and black stars indicate clock- and counterclockwise pinwheels as in **A**.

**Figure 4.**
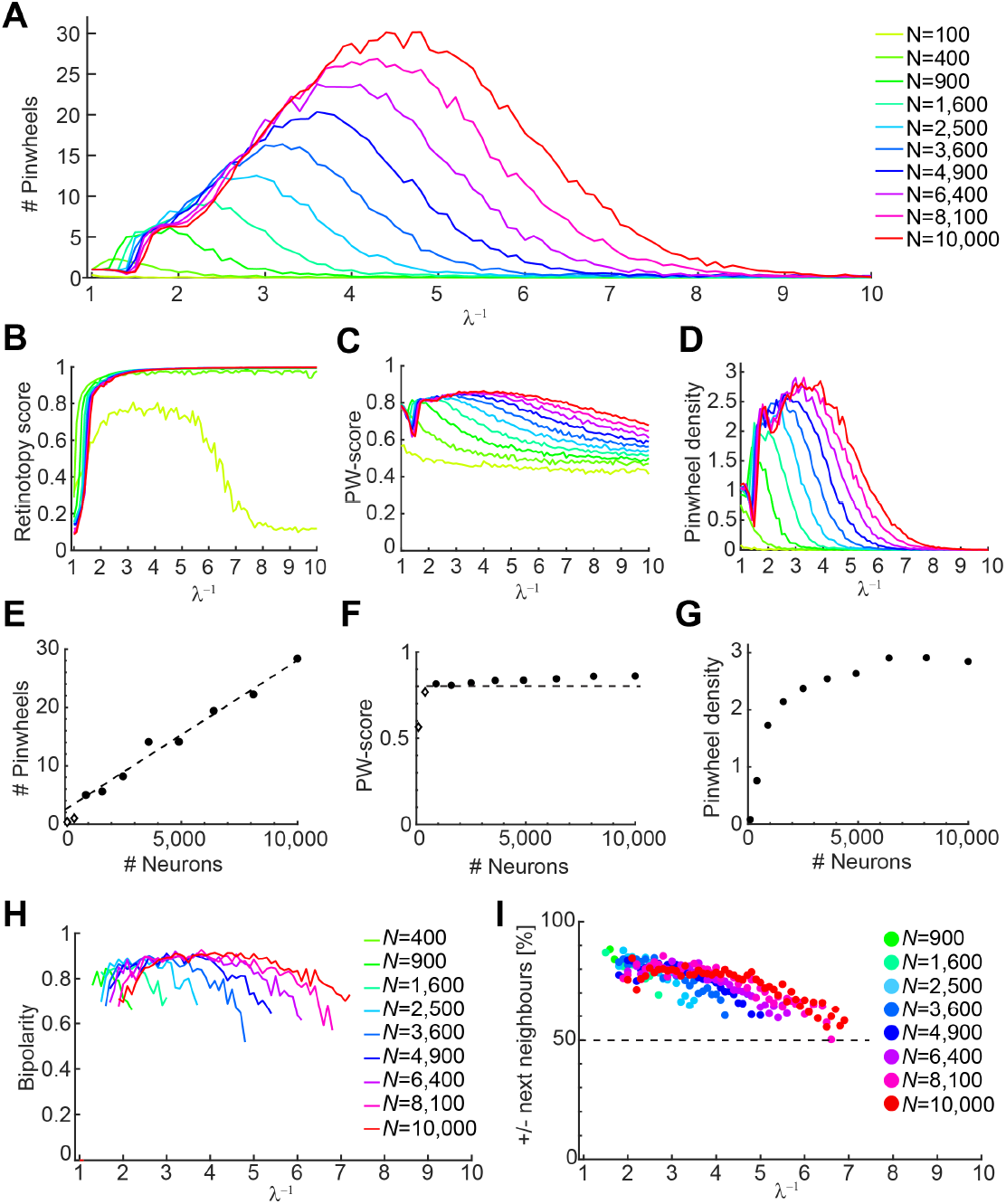
The visual cortex model also indicates a linear relationship between numbers of neurons and pinwheels for similarly structured OP maps. **A**, Number of pinwheels in dependence of receptive field size for different numbers of neurons (see legend). Each of the curves connects 91 data points (from *λ* = 1 to *λ* = 10 in 0.1 steps) where each data point is the mean number of detected pinwheels for 50 trials. **B**, Measured retinotopy scores (see **Materials and Methods**) for the retinotopic maps corresponding to the instances shown in **A. C**, Mean of the maximum PW-score (see **Materials and Methods**) to measure OP map structure for the instances shown in **A**. The bend in the curve between 1 and 2 can be explained by a phase transition from a single to multiple pinwheels in this range of receptive field sizes. For 100 and 400 neurons no more than one pinwheel is observed in most instances as shown in **A** which is why there is no bend in these curves. **D**, Mean normalised pinwheel densities 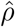 (see **Materials and Methods**) for the instances shown in **A**. The bends in the curves represent the transition from a single to multiple pinwheels as in **C. E**, Numbers of neurons and pinwheels at the points of maximum pinwheel density shown in **D**. Linear fit is indicated by black dashed line (*R*^2^ = 0.9807). Points shown as diamonds were excluded from the fit. **F**, Maximum PW-scores (Mean values for 50 trials) for the instances in **E** as a proxy for the OP map structure. Diamonds indicate the instances that were excluded from the linear fit in **E. G**, Maxima of the mean normalised pinwheel densities 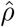 in **D. H**, Mean bipolarity (see **Materials and Methods**) describing the distribution of clock- and counterclockwise singularities. Bipolarities were calculated for every applicable parameter combination of neuronal numbers *N* (indicated by different colours as shown in legend) and receptive field sizes *λ*. **I**, Percent singularities where the nearest neighbouring singularity were of opposite polarity (see **Materials and Methods**) for every applicable parameter combination of neuronal numbers (indicated by different colours as shown in legend) and receptive field sizes *λ*.

The connection probability between neurons in the visual cortex depends also on the OP difference between neurons such that neurons with a more similar OP have a high connection probability and vice versa (Ko et al., 2011, 2013; Martin and Schröder, 2013; Lee et al., 2016). Accordingly, we set the connection probability 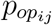 based on the difference between the orientation preferences of neurons *i* and *j* to

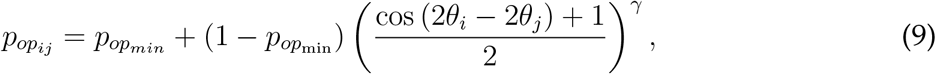

where 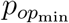 was the minimum connection probability for orientation preference and *γ* the selectivity of the orientation preference. We set 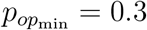 and *γ* = 0.3 for all calculations (instances of connection function shown in **Figure S8A**, *right*). The overall connection probability between neuron *i* and *j* was then given by

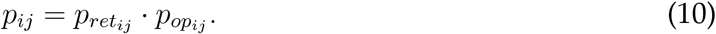

To show that our model was also able to generate realistic maps for more than two features we added the ocular dominance as a third feature to our model neurons. Ocular dominance *ω* was randomly assigned by setting either a 0 for left or a 1 for right eye dominance (*ω* ∈ Ω, Ω = {0, 1}). The connection probability between two neurons based on their ocular dominance was then defined by

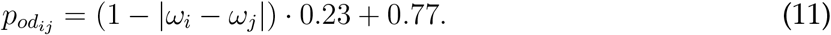

The overall connection probability in that case was

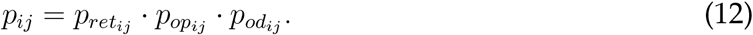

The different feature preferences of the model neurons were visualised by the colour of the dots (**Figures 2C, 3A, 3C and 5**). Colours for orientation preferences were periodic with the hue changing according to the hsv colormap in *Matlab* (**Figures 2C and 5**). Retinotopy was visualised by colours from the viridis colormap. Colours depended here on the retinotopic coordinates *x* and *y* that were visualised separately in paired plots, where purple was the minimum value and yellow the maximum value of the corresponding retinotopic coordinate component (see legend in **Figures 2C and 5**). For visualising the ocular dominance of neurons we indicated left eye dominance by grey and right eye dominance by black (**Figure 5**, *top right*). Distorted maps were also obtained for the visual cortex model and excluded from the analysis (see representative examples in **Figure S2C**).

**Figure 5.**
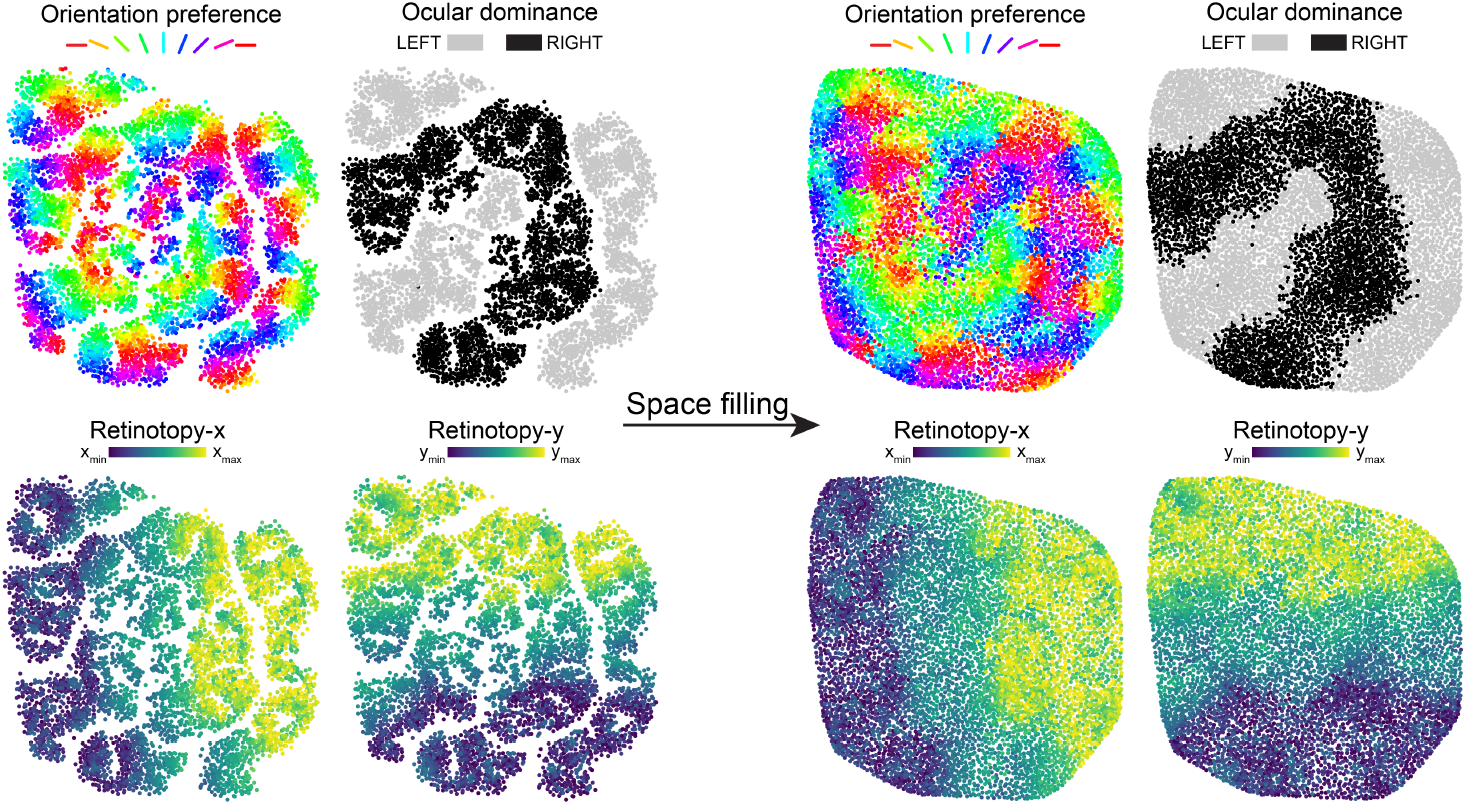
Adding more features to the visual cortex model and space filling. Using a connectivity based on retinotopy, OP and additionally ocular dominance (see **Materials and Methods**) with 6, 400 neurons and *λ*^*-*1^ = 2.2, the resulting OP maps contained pinwheels, the ocular dominance map displayed thick stripes corresponding to either left or right eye preference and the retinotopic preference map showed a gradual shift in the retinotopy of neurons. These maps matched the general phenotype described for example in macaques and cats (Obermayer and Blasdel, 1993; Engelmann et al., 2002). (*left* to *right*) Applying a procedure that promotes space filling in the arrangement leads to more evenly distributed neurons while still preserving the general layout of the map. (*right*) The resulting arrangement after 80 rounds of the space filling procedure (see **Materials and Methods** and **Figure S7**). Feature preferences are indicated by the different colours as shown in the legend for each map.

### Detecting pinwheels

To detect pinwheels in our modeled visual cortex maps, we stochastically sampled a *pinwheelness* score (PW-score). Based on the coverage and continuity of the OP feature space around a sampled location, this score indicated how well this location represented the center of a pinwheel (**Figure 4A**, *middle*). The PW-score was a composite of two separate components. The first component was the correlation to the azimuth (CTA) that has previously been used to quantify the map structure given by the continuity of orientation preferences in a pinwheel (Ohki et al., 2006; Weigand et al., 2017). The CTA measures the Pearson correlation coefficient between the OP of neurons and their angular displacement around the pinwheel center with respect to a reference vector. The reference vector in this case was defined by the mean direction of the neurons that are in the 10^*th*^ percentile of the smallest orientation angles. To quantify the structure of a pinwheel, the CTA needed to be calculated at the pinwheel center. Since the pinwheel centers are per se unknown in our case, we combined the CTA with a second score, which enabled us to predict the pinwheel centers. The second score quantified how well the space of the different OPs was covered in the neighbourhood of each sampled point.

The practical steps of calculating the PW-score consisted of first defining which neurons were in the neighbourhood of the current sample point. In our analysis we defined 20 angle sections that were equally spaced on the full circle around the sample point. For each angle section we then selected the nearest 7 neurons for the further calculations. For those selected neurons, we calculated the CTA as described previously (Weigand et al., 2017). The CTA was a value between *-*1 and 1 where *-*1 represented a perfect counterclockwise pinwheel, 1 a perfect clockwise pinwheel and 0 a perfect salt-and-pepper pattern. The CTA was used to detect pinwheels (**Figure 3A**, *left*), when the values exceeded *>* 0.6 for counterclockwise pinwheels (black stars) or went below < *-*0.6 for clockwise pinwheels (gray stars). Furthermore we introduce a value measuring the coverage of the orientation preference space Θ.

For this purpose, we divided the space of possible orientation preferences into 20 bins of equal sizes. We then checked how many of these bins covered the orientation preference of at least one of the selected neurons. The coverage score was a value between 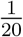 and 1 where 1 indicated a complete coverage of all 20 bins. Both scores were multiplied to obtain the PW-score, a value between *-*1 and 1 where 1 represents a perfect clock- and *-*1 a perfect counterclockwise pinwheel indicated as colours between magenta and cyan at the sampled location (**Figure 3A**). We sampled the PW-score for 20, 000 points to finally detect the pinwheels by two additional steps. First, we selected all sample points with a score above 0.6 (**Figure 3A**, *right*; yellow crosses). Second, we performed a clustering in dependence of sampling point position and its PW-score using the density-based spatial clustering of applications with noise (DBSCAN) algorithm (with parameters *minPts* = 5 and *ε* = 10) (**Figure 3A**, *right*; black dashed lines). The pinwheel centers were defined as the center of mass of the clusters (**Figure 3A**, *right*; red asterisks).

### Calculating retinotopy score

In order to quantify the structure of the retinotopic arrangement in our modeled maps, we considered the correlation of the neuronal coordinates in space and their retinotopic feature preference. However, the correlation varied depending on which axis of the spatial and retinotopic coordinates were used and how the neuronal map was rotated relative to the spatial axes. Therefore, we calculated the correlation for all combinations of axes and rotated the map using a local search algorithm until the best correlation value was found. We took the absolute value of the correlation as the retinotopy score.

### Measuring geometric orientation hypercolumn size and calculating normalised pinwheel density

To calculate the pinwheel density normalised by the orientation hypercolumn size 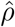 we calculated the latter by first sampling the cosine of the orientation preference at 100 linearly spaced points along a virtual electrode track. The electrode track was defined by a connecting line between a randomly selected neuron and the farthest neuron from the selected neuron. We calculated the power spectrum of the spatial frequency of the sampled values. From the power spectrum we selected the three largest peaks and calculated the weighted mean of the spatial frequency given by the power of the peaks and their frequency. Using the resulting spatial frequency *v* and the length *l* of the electrode track, the orientation hypercolumn size Λ was given by

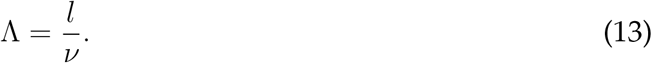

This procedure was repeated for 50 times and Λ was defined by the median of these values. We then calculated the spatial pinwheel density *ρ*:

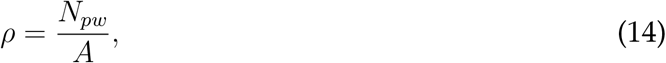

where *A* was the area of the modeled map and *N*_*pw*_ the number of pinwheels. Finally, the overall normalised pinwheel density was then calculated as

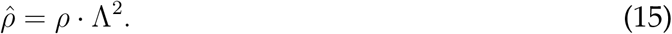

### Statistics of pinwheel singularities

In each OP map, we calculated the bipolarity that was 1 if the number of clockwise singularities matched the number of counterclockwise singularities and 0 if either only clock- or counterclockwise singularities were present (**Figure 4H**):

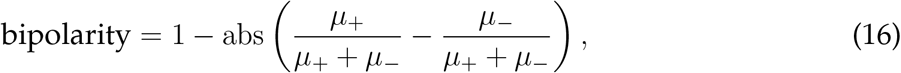

where *µ*_+_ is the number of clockwise and *µ*_*-*_ the number of counterclockwise singularities. Based on this definition the bipolarity is only defined if *µ*_+_ + *µ*_*-*_ *>* 0. Therefore, the mean bipolarity was only calculated if pinwheels were detected in all instances of a certain parameter combination of numbers of neurons *N* and receptive field sizes *λ* (**Figure 4H**). We further calculated how likely it was for nearest neighbour singularities to be of opposite polarity in percent (**Figure 4I**). The relative amount of nearest neighbour singularities was only defined if at least two pinwheels were detected in an OP map and its mean value was only calculated if this condition was fulfilled in all instances of a certain parameter combination of numbers of neurons *N* and receptive field sizes *λ* (**Figure 4I**).

### Sampled visual cortex maps

To test our pinwheel detection method, we sampled OP maps from different mammalian species that were shown in (Kaschube et al., 2010). We enhanced brightness and contrast of the images since colour information was lost due to the usage of CMYK colours in the document. We stochastically sampled the colour at 5, 000 points in these maps (*bottom*). To derive the OPs that were given to the neurons in our model, we had to set the OPs for the sampled points according to their colours. To accomplish this, we created a colormap of 100 equidistant points in the hsv colour space in *Matlab* and assigned each colour one of 100 linearly spaced orientations from 0 to *π*. The OP of each sampled point was then set according to the assigned OP of the colour that had the smallest distance to the colour of the sampled point.

### Space filling

To optimise the space filling of a predicted neuronal arrangement (**Figure 5**), we first generated a 500 × 500 square lattice **L** inside the boundaries of the minimum and maximum coordinates of the positions of the model neurons **Y**. We further discarded the points of **L** that lay outside of the convex hull of **Y**. The remaining grid points **X** of the square lattice **L** were used to detect the density of model neurons in the neuronal arrangement, which allowed for the optimisation of a better space filling by iteratively shifting the positions **Y** towards less dense regions in the arrangement. In each iteration *i* of the algorithm, a grid point **x**_*i*_ was randomly selected by a probability *p* that decreased exponentially with the minimum distance of **x**_*i*_ to any position **x**_*i*_ of the positions **Y**. The model neuron position 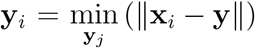 with the minimum distance to the selected grid point **x**_*i*_ was then shifted towards **x**_*i*_ such that **y**_*i*+1_ = **y**_*i*_ + 0.5 *\** (**x**_*i*_ *-* **y**) in iteration *i* + 1 of the algorithm. Each round of this optimisation procedure consisted of *N* iterations and in each round the grid was randomly shifted by a small amount (**X** + *ε* ∈ [0, 1]) to avoid generating a completely regular arrangement. After 80 rounds a relatively even distribution of representative examples was reached that covered the range of the different modeled map phenotypes (**Figure S7**).

### Curated visual cortex data

An overview of the biological data collected for **Figure 1B** is given in **Table 1** and the sources for curated pinwheel density values are given in **Table S1**.

**Table 1.** Biological data. The number of neurons and size of V1 for macaque, owl monkey, galago, squirrel monkey, tree shrew, cat, rabbit, squirrel, rat and mouse were taken as the rounded mean from previously curated data (Weigand et al., 2017). The number of V1 pinwheels was calculated as the rounded values of the multiplication of the curated mean pinwheel densities given in **Table S1** and V1 size. The number of neurons per pinwheel was calculated by dividing the number of neurons by the number of pinwheels in V1 and rounded to the nearest integer. The number of orientation hypercolumns was calculated by 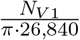, where *N*_*V*1_ is the number of neurons in V1 and *π*·26, 840 the number of neurons per orientation hypercolumn since the number of pinwheels per orientation hypercolumn has previously (Kaschube et al., 2010) been approximated as and the number of neurons per pinwheel is approximately 26, 840 (**Figure 1B**). (PW = pinwheel, HC = orientation hypercolumn). The italic numbers in parentheses for rodent species are estimates if orientation hypercolumns would hypothetically exist in these species. They were calculated based on the determined approximate orientation hypercolumn size of 26, 840 neurons and the number of neurons and size of V1 in the respective species.

| V1 | #Neurons | Size [mm<sup>2</sup>] | #PWmm<sup>2</sup> | #PW | #Neurons/#PW | #HC |
| --- | --- | --- | --- | --- | --- | --- |
| Mouse | 314,983 | 3.879 | 0 (3.093) | 0 (12) | 0 (26,840) | 0 (4) |
| Rat | 715,808 | 8.5 | 0 (3.176) | 0 (27) | 0 (26,840) | 0 (8) |
| Squirrel | 2,694,816 | 32 | 0 (3.125) | 0 (100) | 0 (26,840) | 0 (32) |
| Rabbit | 5,920,000 | 74 | 0 (2.986) | 0 (221) | 0 (26,840) | 0 (70) |
| Tree Shrew | 8,097,600 | 42 | 8.375 | 352 | 23,020 | 96 |
| Cat | 27,061,800 | 345 | 2.816 | 972 | 27,853 | 321 |
| Galago | 41,910,636 | 200 | 6.639 | 1,328 | 31,563 | 497 |
| Owl monkey | 56,565,642 | 301.2 | 7.4 | 2,229 | 25,379 | 671 |
| Squirrel monkey | 174,283,200 | 637 | 11.05 | 7,039 | 24,760 | 2,067 |
| Macaque | 240,435,700 | 1072.5 | 7.94 | 8,516 | 28,235 | 2,851 |

## Results

### Optimal neural placement predicts constant neuron numbers per hypercolumn

In order to better understand the relationship of cortical hypercolumns in brains of varying sizes or with different neuronal densities, we first designed a simple model on the premise that neurons belonging to the same hypercolumn are preferentially connected to one another. We used simple grid-like connectivity matrices neglecting the quality of the features represented by the individual neurons. To randomly set the connection between two neurons associated with their respective hypercolumns, the connection probability decayed exponentially (see **Equation 6**) along the Euclidean distance on the grid (**Figure 1A**, *top row*).

Using this connectivity with our neuronal placement procedure, we obtained maps that varied in the number of neurons per hypercolumn and overall grid sizes (**Figure 1A**, similar results were obtained using oMDS, see **Figure S3**). Interestingly, the formation of clearly discernible hypercolumns depended on the number of neurons inside a hypercolumn but not on the total size of the map in terms of the number of hypercolumns. We introduced the measure *s* (see **Equation 7**), the scatter of the respective maps, that quantifies the prominence of individual hypercolumns based on how many of their neurons intruded other hypercolumns (see **Materials and Methods**). The average *s* for 15 trials in each parameter configuration varied strongly for very small map sizes and thus low overall number of neurons but rapidly reached a steady state regardless of neuron numbers *n* per hypercolumn that we tested (prominent hypercolumns at e.g. *s ≤* 0.5 were only reached if *n >* 40, **Figure S4**). The influence of map size on the structure was only marginal for maps that contained prominent hypercolumns. Thus, our simple model would predict that similarly structured cortical maps with similarly prominent hypercolumns emerge with similar numbers of neurons per hypercolumn.

### Pinwheels with constant neuron numbers are consistent with existing data

To investigate whether real cortical structures operate under similar constraints as in our model we used curated data for overall neuronal numbers in visual cortex V1 (Weigand et al., 2017) and newly curated data for pinwheel densities in the six different mammalian species for which the corresponding measurements exist (**Table S1**), summarised in **Table 1**. Analysing these data, we found that the average number of neurons inside one pinwheel is relatively constant at around 27, 000 neurons per pinwheel (slope of the linear fit in **Figure 1B**), in keeping with previous results (Srinivasan et al., 2015). This compares well with the linear relationship found in our model between the number of neurons and the number of hypercolumns using *s* ≈ 0.5 to produce maps with similar structure (**Figure 1C**; see **Materials and Methods**).

Neuron numbers in the model were restricted to < 10, 000 out of computational reasons making a precise quantitative match impossible between model and biology (compare ranges in **Figures 1B and C**). Additionally, in the biological data, of course, the salt-and-pepper arrangement in small species lacks orientation hypercolumns altogether (**Figure 1B**). Considering that a pinwheel consists of about 27, 000 neurons and even the smallest measured rodent species possess at least 10 times more neurons in the visual cortex, this raises the question why orientation hypercolumns have not been observed in all mammals. Although not backed up by empirical results, it has already been discussed that the emergence of pinwheels in smaller rodents is very unlikely because pinwheels cannot exceed a certain size relative to the total visual cortex (Harris and Mrsic-Flogel, 2013). In the following we demonstrate more clearly why a constant orientation hypercolumn size supports this hypothesis and is accordingly not mutually exclusive with the proposed neuronal number dependent phase transition between unstructured and structured visual cortex maps that we observed previously (Weigand et al., 2017).

### Predicted visual cortex maps crucially depend on receptive field size

The size of the visual cortex varies over several orders of magnitudes between mammalian species (**Table 1**) but the size of a single cortical neuron is comparably much less variable (Kaas, 2000; Herculano-Houzel et al., 2014). This relationship between cortical and neuronal size implies that the range of potential connections between neurons with different retinotopic preferences varies considerably in the visual cortex. Correspondingly, the receptive field sizes depend on the visual cortex size. With high neuron numbers in V1, the retinotopic resolution is high and receptive fields are small (**Figure 2A**, *right*). Conversely, low numbers of neurons lead to a lower retinotopic resolution and larger receptive fields (**Figure 2A**, *left*). Assuming that small rodents would possess few orientation hypercolumns as determined earlier (**Figure 2B**), they would be so large relative to the total V1 area that only one or a few orientations could be represented in any part of the visual field (Harris and Mrsic-Flogel, 2013). Alternatively, the retinotopic arrangement could be distorted in favor of a structured OP map, but this has not been observed in mammals yet (Swindale, 2008). In order to validate this conceptual notion numerically, we implemented a V1 model based on our neuronal placement method. Here, the neuronal connectivity used for the placement was modeled by virtue of specific feature preferences – retinotopy and orientation – that were separately randomly assigned to each neuron. The pairwise similarity of the feature preferences were combined as connection probabilities to randomly connect the modeled neurons (see **Equations 8, 9 and 10**). Analogously to the biological case, the receptive field size *λ* determines the connection probability between model neurons based on their retinotopic preference (see **Equation 8**).

Using our model we were able to qualitatively assess the effects of receptive field sizes *λ* and number of neurons *n* by varying both parameters independently. However, due to computational limitations we were not able to make quantitative predictions since the number of model neurons that we could use was practically bounded above 10, 000 neurons. The receptive field size was varied in our model by changing the decay of connection probability with the difference in retinotopic preference between neurons (see **Equation 8**). Increasing the receptive field size without changing the number of neurons led to larger orientation hypercolumns and a more structured OP map (**Figure 2C**, *top*; right to left). A similar effect was found when decreasing the map size but increasing virtual hypercolumn sizes to keep the total number of neurons fixed in our simple grid model (**Figure S5**). Cortical maps with multiple features were only found using t–SNE (compare **Figure S6** with oMDS). The clear OP maps come at the cost of a compromised retinotopic map that eventually collapses entirely (**Figure 2C**, *bottom*; right to left). Conversely, decreasing receptive field sizes results in collapsing the columnar arrangement of orientation preferences and a salt-and-pepper OP map emerged (**Figure 2C**, *top right*). Thus, to maintain the retinotopic map, an upper bound for receptive field sizes should exist. Such an upper bound could define a critical number of neurons that is necessary for the formation of a structured OP map and could explain why orientation hypercolumns are absent in all yet investigated rodent species (Kaschube, 2014). In the following, the dependency between map structure, receptive field size, number of neurons and pinwheels is systematically analysed.

### Detailed model confirms constant neuron numbers per pinwheel

In order to systematically analyse the dependency between map structure, receptive field size, number of neurons and pinwheels in our visual cortex model, we needed to detect pinwheels automatically in the predicted maps. Pinwheels were detected by sampling a score in the predicted maps that indicated whether a clock- or counterclockwise pinwheel was more likely present at each sampled position (**Figure 3A**, see **Materials and Methods**). After applying a threshold to the sampled scores, the remaining points were potential pinwheel centers (yellow crosses in **Figure 3A**, *right*). Applying the DBSCAN cluster algorithm on this point set delivered separate clusters of potential pinwheel centers and the center of these clusters defined the detected pinwheel centers (red stars in **Figure 3A**, *right*). Using the detection method with neuronal arrangements sampled from images of OP maps from different mammalian species (Kaschube et al., 2010), showed that most of the pinwheels were detected (56 out of 57) and that the respective relative pinwheel densities were in the expected range around *π* (**Figure 3B**).

Based on our visual cortex model, we performed similar analyses as we did for the simple grid model (**Figure 3C**). Similar to **Figures 1 and S3** in which the grid size fixes hypercolumn numbers, receptive field sizes in the V1 model determined the number of pinwheels. Using the maximum of the sampled pinwheel scores as an approximation for the OP map structure (PW-score, indicated by the brightest cyan or magenta coloured dot in **Figures 3A, B**) showed that the number of neurons per pinwheel was constant for similarly structured maps irrespective of receptive field size (**Figure 3C**). This was comparable to the results of the simple grid model where regardless of map size the number of neurons was constant for similarly structured maps (**Figure 1A**).

To analyse this potential constant relationship between the number of neurons and pinwheels numerically, we generated a large dataset of models spanning a wide range of the parameter space (**Figure 4**). In the parameter space combining large receptive fields (*λ*^*-*1^ < 2) and small neuron numbers *N* the existence of both pinwheels (**Figure 4A**) and retinotopy (**Figure 4B**) was not possible. This finding indicates that there exists an absolute upper bound for the receptive field size and therefore a lower bound of neuronal numbers for the emergence of structured OP maps (**Figure 4A, and C**). Interestingly, with a very small number of neurons, even the retinotopic map did not reach the amount of structure found for higher number of neurons and even vanished entirely at small receptive field sizes (**Figure 4B**, *N* = 100). We further determined whether our V1 model showed a linear relationship between the number of pinwheels and neurons. Since a peak of the measured PW-scores was not as clearly defined (**Figure 4C**), we used the peak of the normalised pinwheel density (see **Materials and Methods**) in **Figure 4D** as a reference point that defines a similar map structure. At this point the best compromise between a clearly structured OP-map and a concurrently least distorted retinotopic map is reached, because pinwheels are abundant throughout the map but also as small as possible. The measured relationship between the number of neurons and pinwheels in these similarly structured maps was indeed linear (**Figure 4E**, *R*^2^ = 0.9807). Measures for very low neuron numbers (*N* = 100, *N* = 400) were excluded from the fit since either no pinwheels were observed (**Figure 4A, and E**) or the retinotopic map was strongly distorted (**Figure 4B**) and the Pinwheel score was too low (< 0.8) (**Figure 4F**). However, the linear relationship was still preserved when those measurements were included (*R*^2^ = 0.9801).

### Detailed model is consistent with characteristic properties of V1 maps

Our V1 model is useful to study and better understand the relationships between many further visual cortex map properties. We show that the normalised pinwheel density (**Figure 4D**) converged to a constant number slightly below 3 (**Figure 4G**), which is in the range of the values found previously in a model and in different mammalian species (Kaschube et al., 2010). Corresponding to other biological results (Swindale, 1996; Obermayer and Blasdel, 1997), clock-and counterclockwise pinwheel singularities were equally represented in our maps (**Figure 4H**) and singularities of opposite polarities tended to be neighbours (**Figure 4I**) because they were alternating in a quasiperiodic fashion (**Figure 4A**). With either very large or small receptive field sizes, these relationships faded because the maps consisted of either one large pinwheel or were near salt-and-pepper maps expressing only few small pinwheels (compare **Figure 4A** and **Figure 4H, I**). The relative amount of singularities with nearest neighbours of opposing polarity was 83.17 *±* 3.74% for the instances used in **Figure 4E–G** which closely matched the mean values of 83.17 *±* 4.71% from 12 macaques and 1 squirrel monkey (calculated from Table 3 in Obermayer and Blasdel, 1997). Details such as the very slight increase in map structure with the overall number of neurons observed in **Figure 4F, E** also match experimental data (reviewed in Weigand et al., 2017). As a proof of principle, we show that the model can handle even more features than retinotopy and OP alone. By also including ocular dominance as a feature, the resulting maps show thick ocular dominance bands in the centers of which pinwheel singularities are preferentially located (**Figure 5**) which is remarkably similar to biological observations (Swindale, 1996; Obermayer and Blasdel, 1997). However, in all of our predicted maps, neurons tended to be distributed irregularly. While the distribution of neurons in the visual cortex is not strictly regular (Ohki et al., 2006), many of our predicted arrangements were obviously much more irregular and contained occasional gaps (**Figures 2C and 3C**). Although the general map layout was not affected by this irregular distribution, we implemented an additional procedure that modified the predicted arrangements for a better space filling. Applying for example this procedure to the neuronal arrangement in **Figure 5** (*left*) led to more homogeneously distributed model neurons while preserving the general appearance of the maps (**Figure 5**, *right*). The corresponding code is available in our model toolbox and allows *post hoc* modifications of any model maps.

## Discussion

Using a novel neuronal placement model, which is loosely based on the premise that wiring length has to be optimised, we showed in accordance with newly curated biological data that orientation hypercolumns in the primary visual cortex appear at fixed neuronal numbers. Given that the number of neurons per orientation hypercolumn is constant, the size of a pinwheel can only vary with neuronal density. We propose that variations in the spatial density of pinwheels between mammalian species could be a mere consequence of similar OP maps that are differently scaled versions of each other. Our results are corroborated by the model’s faithful replication of a multitude of characteristic properties of visual cortex maps.

### Scaling behaviour in OP maps

It is interesting to study the particular scaling behaviour of homologous biological structures (Schmidt-Nielsen, 1984; Kaas, 2000) to better understand their function. The area of the primary visual cortex in different mammalian species varies over several orders of magnitude (Kaas, 2000), which is less a consequence of differences in neuronal density than it is a result of different neuronal numbers in this area (**Table 1**). Depending on visual cortex size, the potential difference in the retinotopic preference of a neuron to its connected neighbours varies (Elston et al., 1996; Kaas, 2000) because the receptive field *λ* of neurons is inversely correlated with the number of neurons in the visual cortex (**Figure 2A**). In our visual cortex model, we found that for smaller numbers of neurons the range of *λ* where orientation hypercolumns are present is smaller. At very low numbers of neurons, orientation hypercolumns do not appear at all. From our curated biological data we propose a lower limit of ∼300 pinwheels in V1 considering that the lowest number of pinwheels is 352 in tree shrew and that rabbits could hypothetically host 221 pinwheels but they do not (**Table 1**). This lower bound presumably exists for the same reason as in our model. With low overall numbers of neurons in V1 the receptive field size of neurons increases but the number of potential connections to neurons of similar OP is too small for pinwheels to form without distorting the retinotopic map. However, with increasing numbers of neurons in V1, pinwheels start to emerge because the local change in retinotopic preference gets small enough such that sufficient connections between neurons of similar OP can form which enables them to cluster and concurrently maintain the retinotopic map. Accordingly, a selective patchy connectivity (Gilbert and Wiesel, 1989; Malach et al., 1993; Fitzpatrick, 1996; Bosking et al., 1997; Lund et al., 2003) is found in species with pinwheels and a local (Van Hooser et al., 2006) but selective (Ko et al., 2011, 2013, 2014; Lee et al., 2016) connectivity is observed in rodents. If recruiting neurons for the formation of a pinwheel was mainly limited by their retinotopic difference, the number of neurons per orientation hypercolumn should steadily grow with increasingly larger visual cortices. However, the number of neurons per pinwheel is constant at around 27, 000 neurons for over one order of magnitude of neuronal numbers in V1. This could be caused by the theoretical limit of on the order of ∼10^4^ neurons for which a potential all-to-all connectivity can exist (Wen and Chklovskii, 2005).

Recently, it has been proposed that different geometric orientation hypercolumn sizes in the visual cortex correspond to the radii of astrocytes (Philips et al., 2017). A dependency of astrocyte radii and orientation hypercolumn size would only be compatible with our results if astrocyte size inversely depended on the neuronal density. Otherwise, the number of neurons per pinwheel would vary strongly between different species, which contradicts our findings. However, for the data given in Philips et al. (2017), astrocyte size does not depend on neuronal density. Therefore, a fixed orientation hypercolumn size could never be maintained. For example the neuronal density in cats is roughly similar to or even higher than that in rodent species (**Table 1**) but the astrocyte size is triple the size given in Philips et al. (2017). In other studies, it has been suggested that the average glial cell size varies only modestly and non-systematically across brain structures and species (Herculano-Houzel, 2011, 2014; Herculano-Houzel et al., 2014). Hence, it remains unknown whether astrocyte size indeed has an influence on orientation hypercolumn size.

### Qualitative predictions from our neuronal placement model

While our models provide qualitative explanations of various biological observations in visual cortex maps, they do not estimate the numbers of neurons quantitatively (**Figure 4A, Table 1**). Given the limits of current computational resources, our models provide good results for neural placements involving up to 10, 000 neurons. Here, we specifically set the parameters such that orientation hypercolumns appeared with low numbers of neurons. In this way, we could analyse a broad spectrum of qualitative features of the resulting maps (**Figures 3 and 4**). Although we specifically selected a range of parameters that enabled us to see salt-and-pepper maps up to maps with a multitude of pinwheels, our model was not very sensitive to parameter changes that affected the connection functions of the visual cortex model (**Figure S8A**). The model nearly always produced maps that corresponded either to a salt-and-pepper or structured OP map even when the otherwise fixed parameters 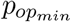 (**Figure S8B**) and *γ* (**Figure S8C**) were varied. Only by using very high *γ* values we could obtain a map pattern that markedly differed from those found in the visual cortex (**Figure S8C**, *rightmost*). Here, individual clusters of neurons with very similar OPs appeared. In that particular case, the tight connection probability around any given orientation preference seemed to force neurons with a very similar OP and retinotopic preferences to be near each other (**Figure S8A**, *right*). Interestingly, some parameters in our model can counteract the effect of other parameters. While high *γ* values tended to result in maps with larger but fewer pinwheels, lowering the receptive field size of neurons led again to maps with more but smaller pinwheels (**Figure S8D**). This indicates that similar map phenotypes could also be produced with different variations of a common connectivity based on the similarity of neuronal feature preferences. It might be possible that with much larger resources, calculations with neuronal numbers in the biological range would become feasible. Accordingly, it would then become possible to characterise the differences in visual cortex maps of different mammalian species quantitatively.

### Comparison between oMDS and t–SNE for optimal neuronal placement

The neuronal arrangements predicted by oMDS and t–SNE can be very similar (compare **Figure 1** and **Figure S3**), but they can also differ markedly (compare **Figure 2C** and **Figure S6A**). In particular, we found that oMDS failed to reproduce visual cortex maps that were based on multiple features. While we could find arrangements that either respected OP or retinotopy, it was never possible to find any that respected both features in the maps obtained using oMDS (**Figures S6A, B**). In comparison to other dimension reduction methods, t–SNE has the advantage that it is able to reveal the structure of high dimensional data at many different scales in the projected space (van der Maaten and Hinton, 2008). That difference to oMDS is probably the reason why t–SNE is able to predict cortical maps based on multiple features. In contrast, oMDS favors arrangements that respect the global rather than the local scale. According to the error function of oMDS, relatively equal deviations between spatial distance in the projected space and transformed connection dissimilarities have a lower impact on the local scale. This is because the absolute differences between small connection dissimilarities and spatial distances are smaller than for larger dissimilarities and distances. Hence, oMDS tends to appropriately map the features that are dominant at the global scale while missing those that could get dominant at the local scale. Since always one of the neuronal feature preferences is at least slightly more dominantly encoded in the neuronal connectivity, oMDS seems to always find the appropriate arrangement for that particular feature ignoring the other feature preferences. It is difficult to objectively measure whether the more realistic predictions by t–SNE compared to oMDS might also be the reason of a superior optimisation of wiring length. Simply measuring the distances between the connected neurons in the predicted arrangements is not a useful comparison because the model neurons are strongly biased towards the middle when using oMDS for the same parameters as used with t–SNE (**Figure S6A**). While such an arrangement trivially leads to a reduction of the summed wiring length by overcrowding the space, overcrowding is not a biologically realistic solution for minimising the wiring length. Therefore, a fair measurement would have to take into account the local densities of neurons throughout the predicted arrangements. Although t–SNE finally enabled us to predict neuronal maps with multiple feature preferences, degenerate solutions appeared regularly that likely corresponded to the optimisation procedure remaining stuck in a local minimum (**Figure S2C**). In general, t–SNE is well known to introduce some structural anomalies when fine-tuning the resulting visualisations, side effects that we have to accept when using our model and interpreting the results (Wattenberg et al., 2016). For example, neurons in our cortical map models were less evenly distributed than in biology (Ohki et al., 2006). Therefore, we do not claim that the precise local spatial relation between neighbouring neurons is biologically realistic whereas the robust solutions obtained for the overall map structures would not be affected by the irregularities occurring at a rather finer level. To obtain a more regular arrangement we also implemented a procedure that increases the space filling in the arrangement (**Figure 5**). However, this space filling procedure comes at a high computational cost. Furthermore, the potential gain in the predicted map structure obtained by this procedure is not obvious, since the arrangements are exclusively modified for space filling without concurrently observing a constraint, which minimises wiring length. Future models could bridge this gap by predicting arrangements that optimise both wiring length and space filling.

### Comparison with other visual cortex map models

Visual cortex models based on elastic net (EN) algorithms are related to our model, since they are also based on dimension reduction. However, there are significant differences. Most importantly, our model changes the positions of model neurons by minimising the Kullback-Leibler divergence of conditional probabilities based on the connection dissimilarity and distance of the model neurons. In contrast, EN models map a multidimensional feature space onto a two dimensional sheet represented by model neurons on a grid. In this case, neuronal positions are fixed while their feature preferences are changed; this happens in the higher dimensional space. The 2*D* sheet of model neurons is folded in the multidimensional space such that a given range of possible feature preferences are visited while the size of the sheet is minimised (Swindale, 1992). The coordinates in the multidimensional space then define the neuron’s respective feature preferences. Accordingly, the EN model effectively changes the feature preferences of model neurons but fixes their positions while in our model the positions of the neurons are changed but the neuronal feature preferences remain fixed.

Kohonen map models also tune the preferred orientation of model neurons fixed on a two-dimensional grid but are not based on a dimension reduction approach. These models instead use a competitive Hebbian learning rule that changes the preferred orientation of model neurons during a learning procedure to predict visual cortex maps. Interestingly, all these models share similar properties with our model and with each other. We already discussed previously that the phase transition in Kohonen map and EN models is reminiscent of a phase transition based on the number of neurons in our model (Weigand et al., 2017). In Kohonen map models a neighbourhood size (Kohonen, 1982; Obermayer et al., 1992) and in EN models a receptive field size parameter (Durbin and Mitchison, 1990; Swindale, 1992; Goodhill and Cimponeriu, 2000) determines the structure of the resulting map. Both, neighbourhood and receptive field size determine the size of a putative area on the grid that is qualified for the formation of a pinwheel, which in principle conforms to the number of neurons that can take part in the formation of a pinwheel. Thus, the phase transition seen in these models is indirectly related to the phase transition depending on neuron numbers. In this regard, the visual cortex model presented here showed in greater detail how the phase transition depends on the size relationships between the number of neurons, receptive field and orientation hypercolumn size (**Figures 2–4**).

We found with our model that in the regime of low neuronal numbers, continuity of the feature space could not be obtained concurrently in both the retinotopic and the OP map (**Figures 4A–C**). Although a continuous representation of the neuronal feature preferences in both feature maps might still be obtainable if the feature space would not be covered at every location of the visual field, it makes no sense to be unable to see some orientations at any specific point of the visual field (Harris and Mrsic-Flogel, 2013). Accordingly, coverage of the feature space even seems to be optimised in the visual cortex of mammals (Swindale et al., 2000). Since possessing both a high coverage and continuity of all feature dimensions is impossible if neuronal numbers are too low, this relationship could explain why structured visual cortex maps are not present in rodents (Weigand et al., 2017). The proposed constant orientation hypercolumn size also supports this hypothesis because it would only allow for a few hypercolumns in small rodents that would inevitably distort the retinotopic map strongly due to their large size relative to the total visual cortex (**Figure 2B**).

Optimising coverage and continuity is also a unifying characteristic of the different models of visual cortex maps (Swindale, 1996; Goodhill and Sejnowski, 1997). While in most models these constraints are a consequence of how the feature preferences of neurons are optimised, our model implements coverage of the feature space by explicitly setting the feature preferences of model neurons such that uniform coverage is ensured (see **Materials and Methods**). However, some studies argue against visual cortex maps being a result of optimising coverage and continuity (Carreira-Perpiñán and Goodhill, 2002; Keil and Wolf, 2011). For example, maps produced by EN models appear to be realistic only if the optimisation procedure is stopped after a certain number of iterations (Keil and Wolf, 2011). This means that results of these models did not correspond to minima in the optimisation procedure. For optimal solutions, realistic pinwheel densities appeared only in extreme ranges of the EN model parameters (Keil and Wolf, 2011). In contrast to this, the maps produced by our model correspond to stable solutions that do not change with a higher amount of iterations (**Figure S9A**). Furthermore, much of the observed variation in the results of our model can be attributed to the variability of the randomly determined connectivities, indicating that our model indeed finds specific solutions for specific connectivities (**Figure S9B**). Accordingly, our model seems to conform to the hypothesis that visual cortex maps optimise coverage and continuity. Future studies could analyse how important a uniform coverage of the feature space is for the formation of a structured visual cortex map and how a biased distribution of preferred features might affect visual cortex maps in general. This is particularly interesting, since our model enables the prediction of maps for arbitrary distributions of neuronal feature preferences.

## Conclusions

For all of our visual cortex model instances we used the same relative connectivity and only varied the receptive field size and the number of neurons (see **Materials and Methods**). Still we were able to see different map phenotypes in our visual cortex model, namely salt-and-pepper and pinwheel patterns (**Figure 2C**), and a linear dependence between the number of neurons and the number of pinwheels, indicating a constant number of neurons per orientation hypercolumn (**Figure 4E**). Hereby our results render these different aspects of OP maps compatible within a single architectural framework using a unified connectivity. We therefore conclude that visual cortex maps and orientation hypercolumns may be rather homologous structures based on a common design principle despite their different phenotypes across mammalian species.

## Declarations

### Funding

This work was supported by a BMBF grant (No. 01GQ1406 – Bernstein Award 2013), and a DFG grant (CU 217/2-1). The authors declare to have no competing financial interests.

## Acknowledgments

We would like to thank L. Deters, J. Muellerleile and M. Schölvinck for useful comments on the manuscript.

## Author contributions

M.W. and H.C. designed the study and designed the model. M.W. performed the simulations and analysed the data. M.W. and H.C. wrote the paper.

## Supporting information

**Figure S1.**
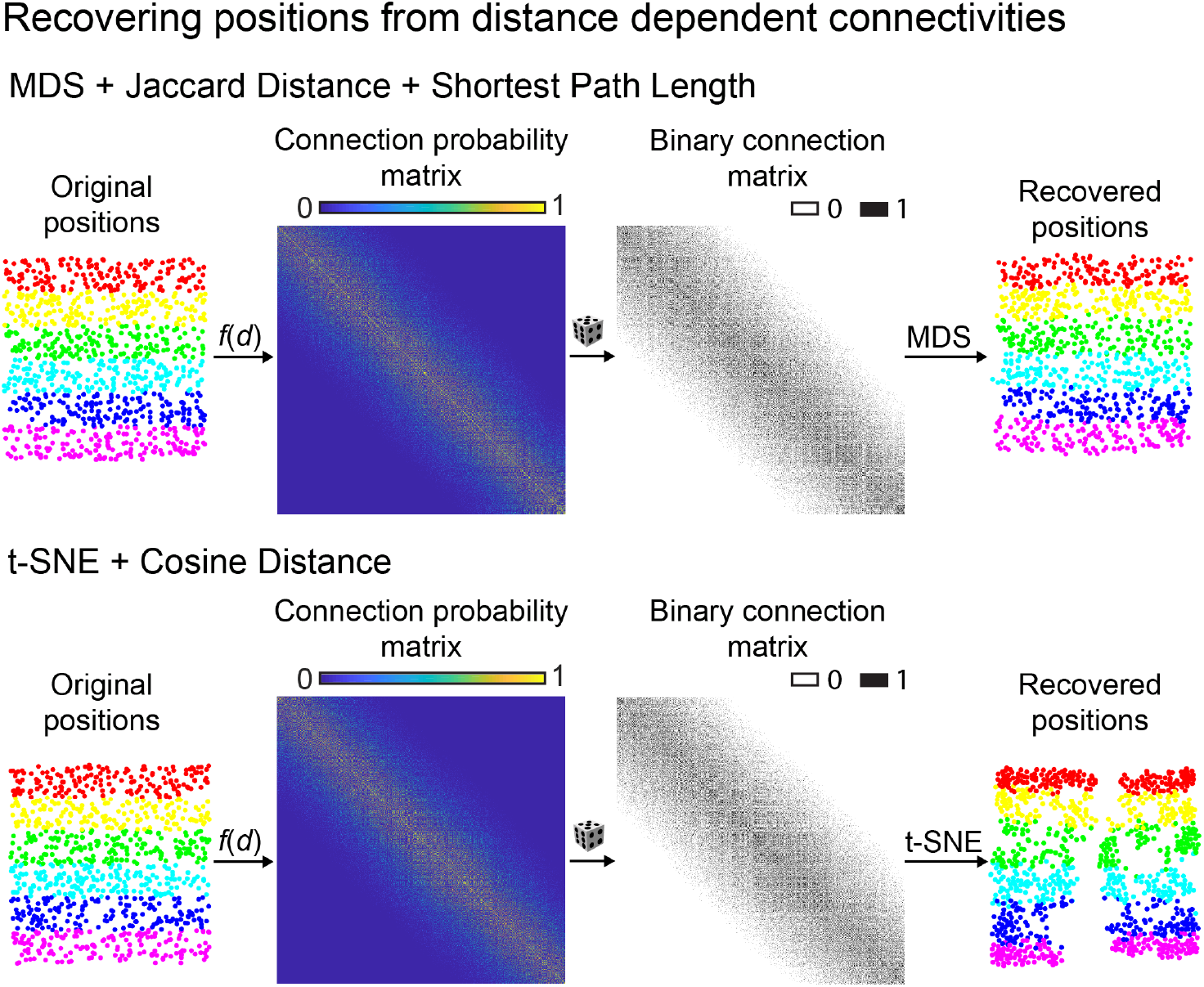
Comparison of oMDS and t–SNE. Recovering random positions divided into six layers (*left*) of equal size by using oMDS with connection dissimilarities obtained by a combination of Jaccard distance (JD) and shortest path length (SPL, *top*, as done in Weigand et al., 2017) and by using t–SNE with connection dissimilarities obtained by cosine distance (CD) (*bottom*). The neuronal connections (*middle right*) are set randomly by the given connection probabilities (*middle left*) which depend on the pairwise distances between points. The recovered positions are shown on the *right*.

**Figure S2.**
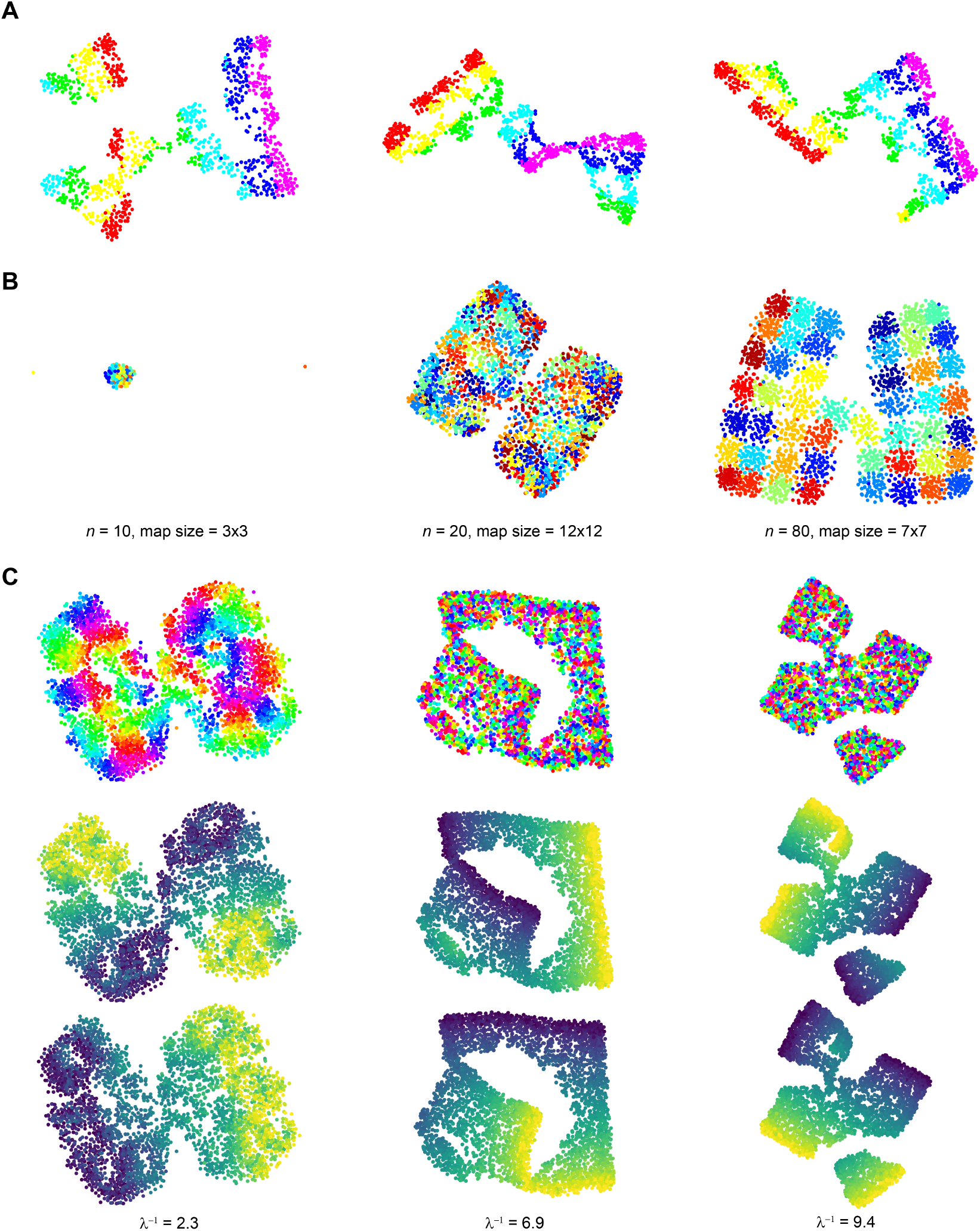
Sample degenerate maps. Representative examples of degenerate maps for **A**, the test setting in **Figure S1, B**, the virtual hypercolumn model in **Figures 1 and S3** and **C**, the visual cortex model in **Figures 2–4**. These degenerate solutions appeared regularly when using t–SNE and were excluded from the analysis. Although distorted maps could be observed for relatively small maps containing fewer amounts of model neurons, the emergence of distorted maps was greatly increased for larger maps containing a relatively high number of neurons likely because increasing the input size led to more local minima in the cost function where the optimisation procedure was trapped.

**Figure S3.**
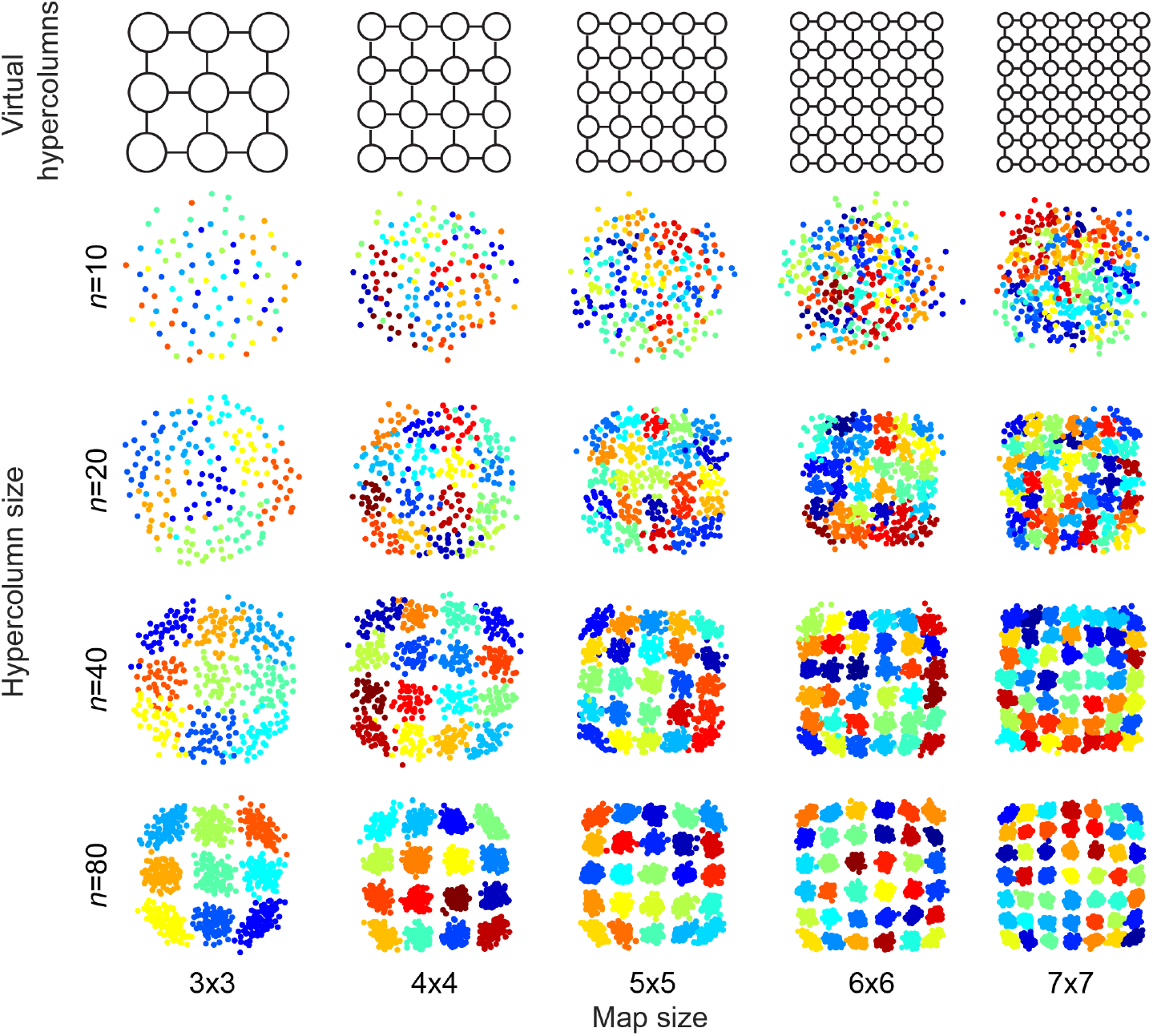
Virtual hypercolumn model results with oMDS as dimension reduction method. As in **Figure 1A**, individual maps of different sizes (horizontal) and different numbers *n* of neurons per hypercolumn (vertical) are shown. Neurons (dots in the maps) assigned to the same hypercolumn in the underlying connectivity matrix are assigned to one random colour.

**Figure S4.**
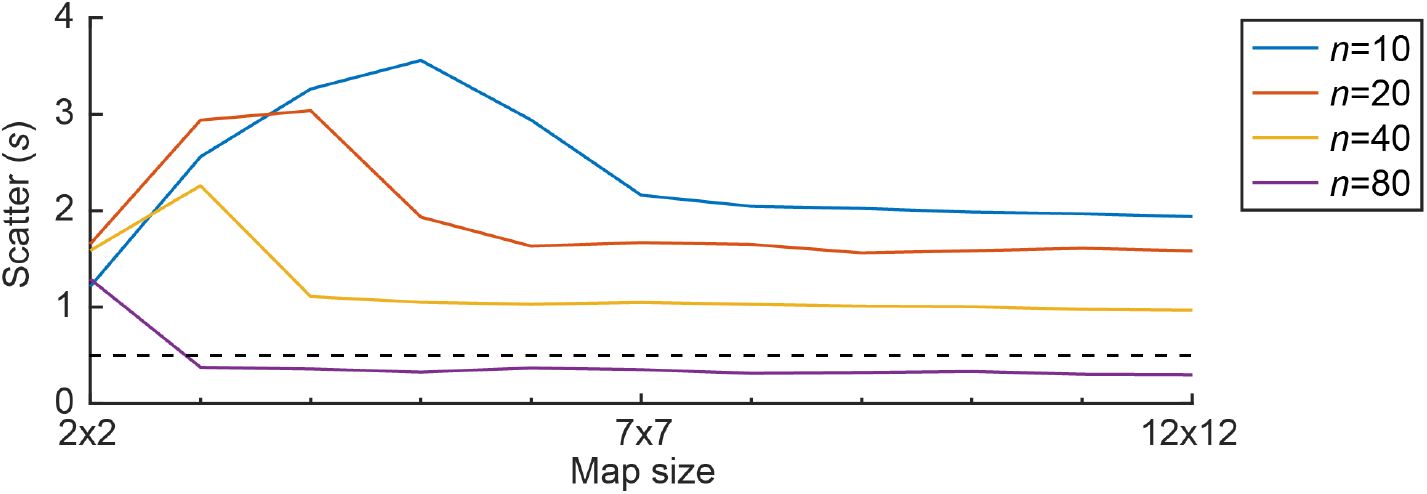
Quantified map structure of virtual hypercolumn model results. Average map structure for different hypercolumn (*n*) and map sizes (15 trials for each parameter combination) quantified by the scatter value *s* (see **Material and Methods**). Dashed line indicates representative threshold value used in **Figure 1C** to define clearly structured hypercolumns.

**Figure S5.**
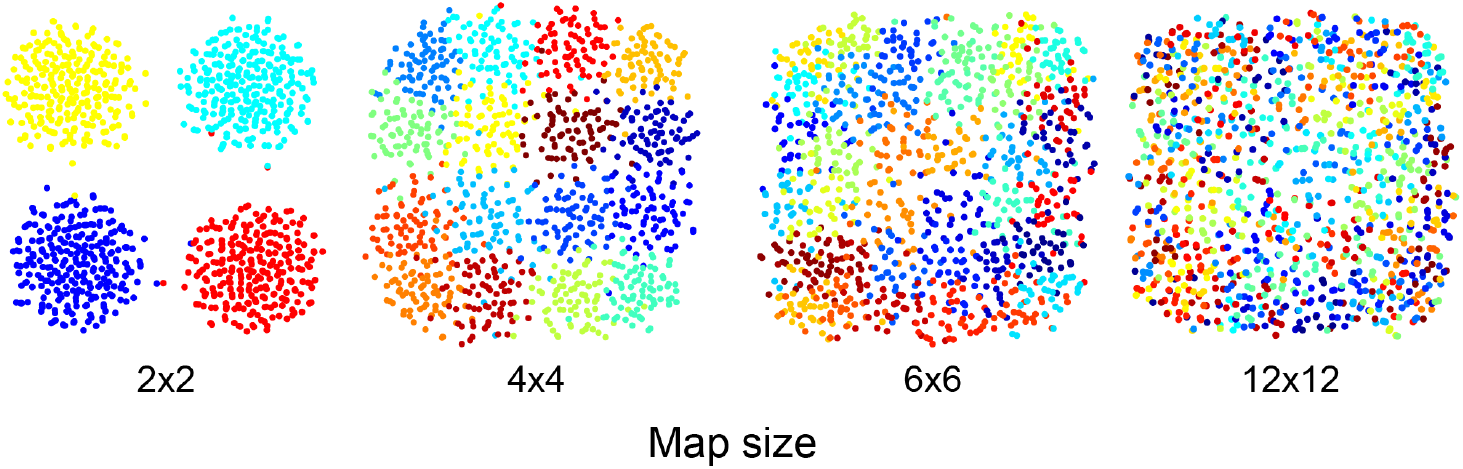
Clamping total neuronal numbers in the grid model. Four maps generated using the same total number of neurons but varying map sizes. To obtain equal total number of neurons we used *n* = 256 for a grid size of 2 × 2, *n* = 72 for 4 × 4, *n* = 32 for 6 × 6, and *n* = 8 for 12 × 12. Increasing the number of neurons per hypercolumn *n* by reducing the number of hypercolumns led to more structured maps.

**Figure S6.**
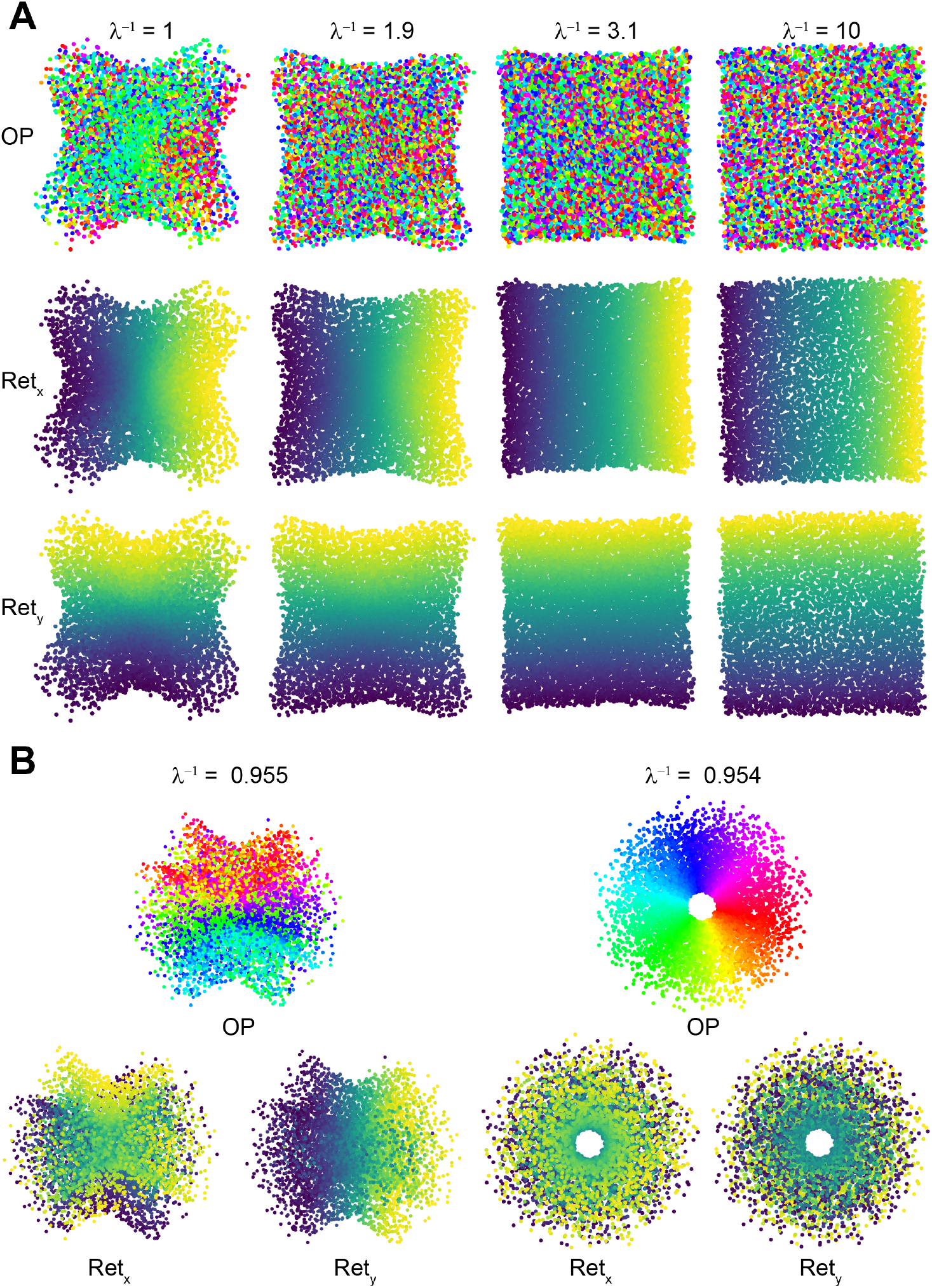
Visual cortex model results using oMDS as dimension reduction method. **A**, As in **Figure 2C**, OP maps (*top*) and retinotopic preference maps (*bottom*) were predicted but oMDS was used instead of t–SNE. Pinwheels did not emerge in this case. **B**, (*right*) Ordered OP map arrangements consisting of one pinwheel can still be obtained when an even lower receptive field size (compared to **A**)) is used. However, this comes at the cost of a structured retinotopic map. (*left*) Concurrently obtaining structured arrangements of the OP and retinotopic map seems to be impossible judging by the sharp transition (when changing *λ*) from a structured retinotopic map to a structured OP map. The number of neurons in all maps was 6, 400 and only the receptive field size *λ* of neurons was varied as indicated. See **Figure 2C** for a legend of the neuronal feature preferences.

**Figure S7.**
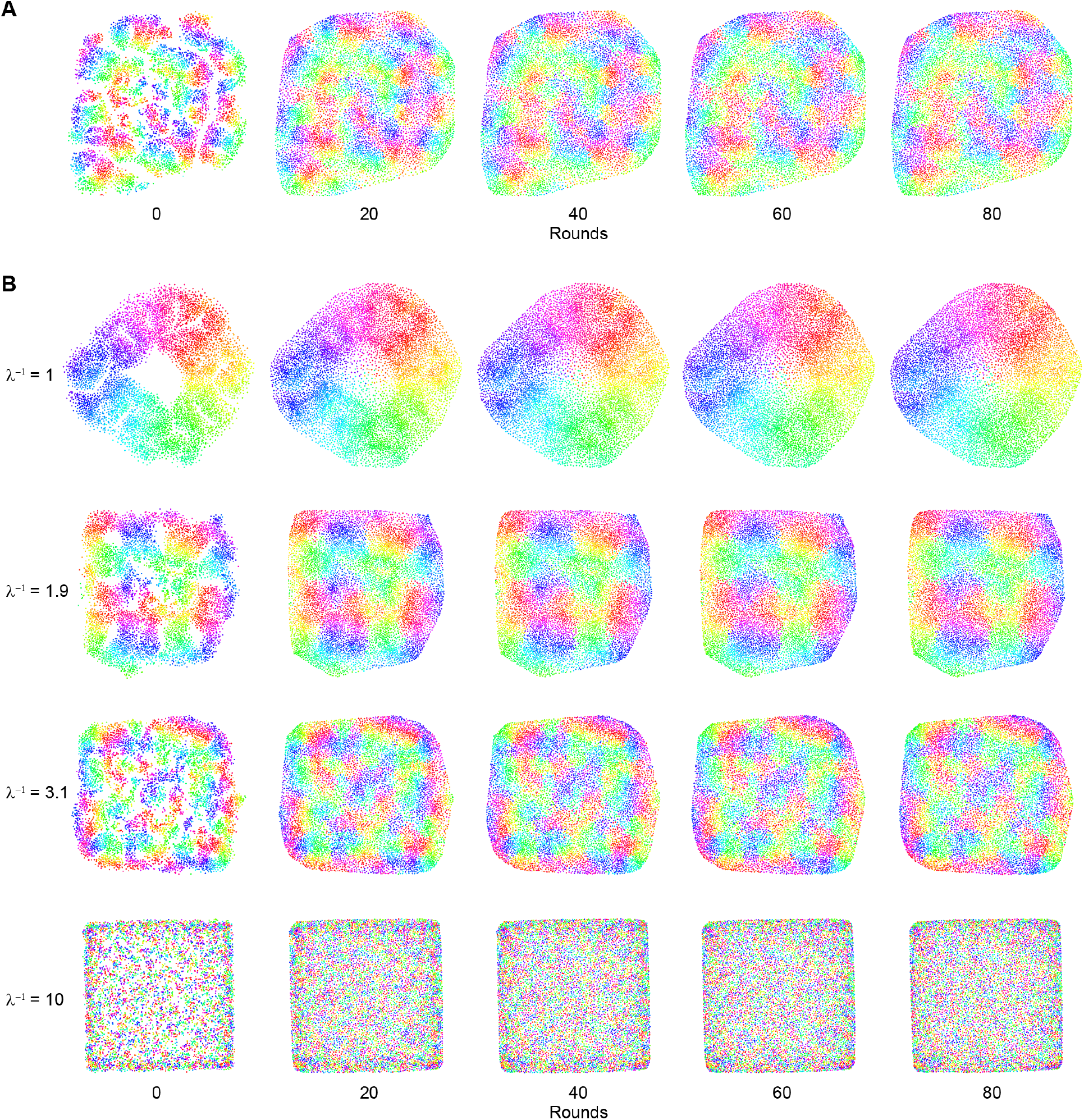
Generating more uniform neuronal arrangements by using the space-filling algorithm. **A**, Progress of the space-filling procedure after different numbers of rounds (see **Materials and Methods**) of the final space-filling result (80 rounds) shown in **Figure 5. B**, Space-filling applied to neuronal arrangements from **Figure 2C** for different values of *λ*. A relatively uniform distribution of neurons is reached after 80 iterations for a wide range of different map layouts with varying *λ*. See **Figure 2C** for a legend of the neuronal feature preferences.

**Figure S8.**
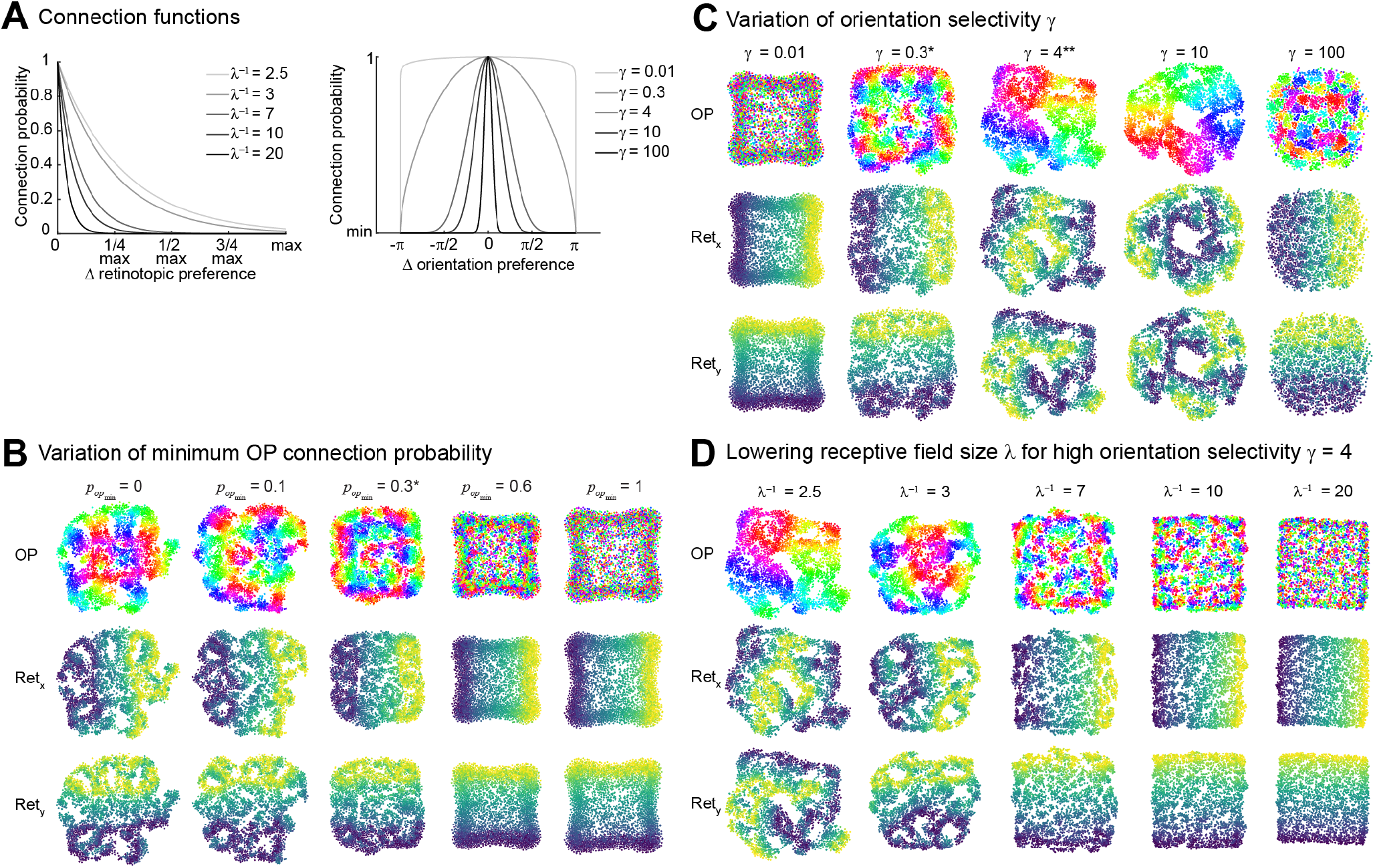
Dependency of visual cortex model on non-varied parameters. **A**, Functions that determine the connection probability based on the pairwise retinotopic and orientation preference difference of neurons. Different connection functions for the varied parameters used in **B, C** and **D** are shown in different shades of grey (see legends). **B**, Different maps for a varied minimum connection probability 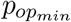 and **C**, a varied orientation selectivity *γ* using a receptive field size *λ*^*-*1^ = 2.5, number of neurons *N* = 3, 600 and otherwise fixed parameters as we used in all other shown results. **D**, Parameters can counteract each other which is shown as an example for the high connection selectivity *γ* = 4 from **C**. Lowering the receptive field size leads again to the formation of multiple pinwheels as for a lower *γ* values in **C**. When using this high connection selectivity, pinwheels can be observed for receptive field sizes that would otherwise be clearly in the salt-and-pepper range (compare with **Figure 3B**). Stars indicate the parameter values that were generally used to obtain all other results. See **Figure 2C** for a legend of the neuronal feature preferences.

**Figure S9.**
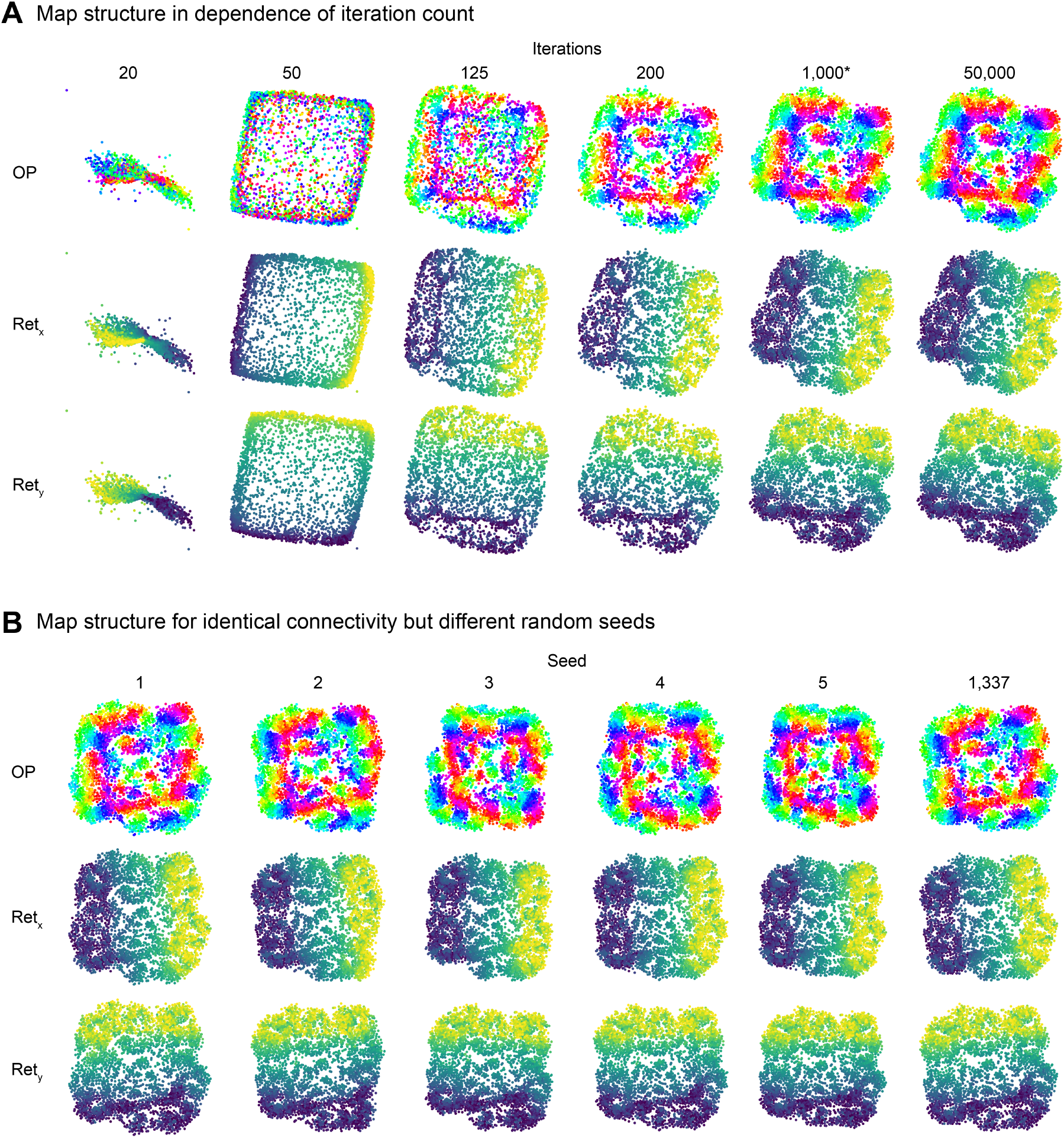
Dependency of visual cortex model on the number of iterations and randomness of the optimisation procedure t–SNE. **A**, Resulting maps for the same random seed but using a different number of iterations during the t–SNE optimisation procedure. After 200 iterations the solution already roughly corresponds to the solution after 1, 000 iterations. Even after 50, 000 iterations no differences to the solution at 1, 000 iterations are recognisable. **B**, Map results for different random seed but the same connection matrix. Recognisable differences between the maps calculated with different seeds exist but the general layout of the map is conserved. Stars indicate the parameter values that were generally used to obtain all other results. See **Figure 2C** for a legend of the neuronal feature preferences.

**Table S1.**
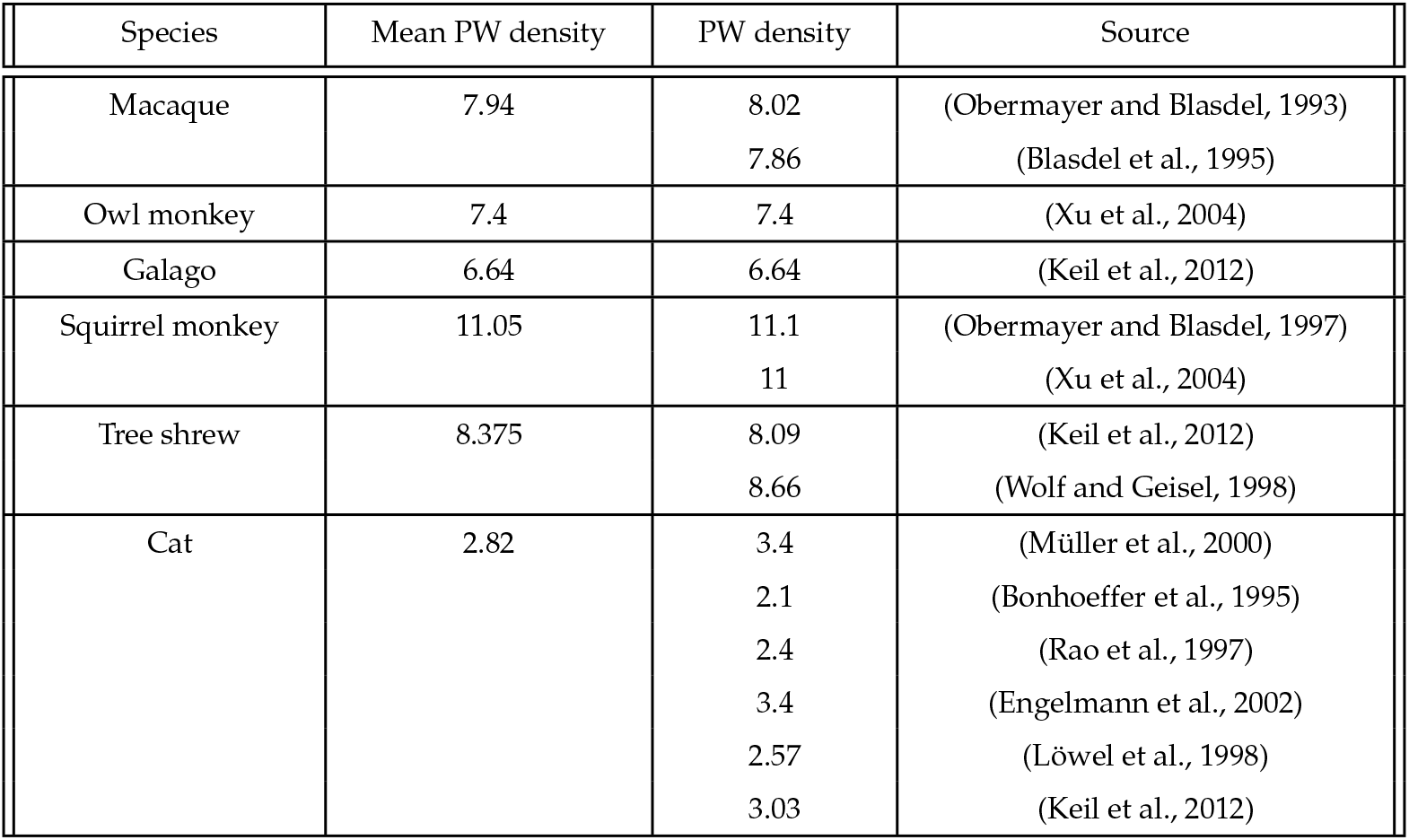
Curated data and sources for pinwheel density values.

| Species | Mean PW density | PW density | Source |
| --- | --- | --- | --- |
| Macaque | 7.94 | 8.02 | (Obermayer and Blasdel, 1993) |
|  |  | 7.86 | (Blasdel et al., 1995) |
| Owl monkey | 7.4 | 7.4 | (Xu et al., 2004) |
| Galago | 6.64 | 6.64 | (Keil et al., 2012) |
| Squirrel monkey | 11.05 | 11.1 | (Obermayer and Blasdel, 1997) |
|  |  | 11 | (Xu et al., 2004) |
| Tree shrew | 8.375 | 8.09 | (Keil et al., 2012) |
|  |  | 8.66 | (Wolf and Geisel, 1998) |
| Cat | 2.82 | 3.4 | (Müller et al., 2000) |
|  |  | 2.1 | (Bonhoeffer et al., 1995) |
|  |  | 2.4 | (Rao et al., 1997) |
|  |  | 3.4 | (Engelmann et al., 2002) |
|  |  | 2.57 | (Löwel et al., 1998) |
|  |  | 3.03 | (Keil et al., 2012) |

## Notes

### Competing Interest Statement

The authors have declared no competing interest.

